# Sensorimotor remapping drives task specialization in prefrontal cortex

**DOI:** 10.1101/2025.09.22.677699

**Authors:** Hugo Tissot, Jeffrey Boucher, Sandra Reinert, Pieter M. Goltstein, Yves Boubenec

**Author notes:** Contributing authors.

## Abstract

The prefrontal cortex (PFC) plays a key role in flexible, context-dependent decision-making. Yet, the population-level mechanisms that support this flexibility remain unclear: is there a generalist neural population involved in representing useful information across tasks, or are representations distributed over multiple modules of task-specialized subpopulations? Here, we investigate whether the population structure modularity in medial prefrontal cortex (mPFC) increases with sensorimotor remapping between task rules in two different paradigms. In a paradigm where all stimuli were remapped, we found a highly modular structure with two task-specialized subpopulations and one generalist. In a different paradigm where only half of the stimuli were remapped, modularity decreased but remained significant. Based on predictions drawn from a recurrent neural network fitted to experimental data, we propose that a gain modulation by global inhibition of mPFC can explain how task-specialized neurons emerge to varying degrees in each paradigm.

## 1 Introduction

As the world changes, we are frequently required to rethink the way we interact with our environment. For instance, the same sensory scene can elicit markedly different behavioral responses depending on the context, which encompasses goals, rules linking stimuli to actions, expectations about forthcoming events, and relevant past experiences [1, 2]. This cognitive flexibility is crucial for animals, allowing them to constantly learn and expand their range of action. Nevertheless, the mechanisms behind this ability are still poorly understood and remain a fundamental question in neuroscience.

Cognitive flexibility is most often studied by training an animal or a neural network on a set of tasks which requires different stimulus-to-output associations. Learning to switch between tasks can range from relatively straightforward to highly challenging, depending on whether sensorimotor mappings are non-conflicting or conflicting across tasks. In a non-conflicting case, different stimuli are mapped to the same output depending on the task (“many-to-one” paradigms, Fig. 1a). Conversely, in other paradigms the same stimuli are mapped to different outputs depending on the task, leading to conflicting associations and requiring sensorimotor remapping (“one-to-many” paradigms, Fig. 1a). Naturally, paradigms can fall between these two extremes and require a mixture of these two components. While these two types of flexibility may appear symmetrical, they impose very different computational constraints. In a many-to-one paradigm, one can find a function *f* which performs the two mappings without having to process the task identity. On the contrary, in a one-to-many paradigm, there is no single function *f* that can perform the two mappings without integrating task information.

**Fig. 1.**
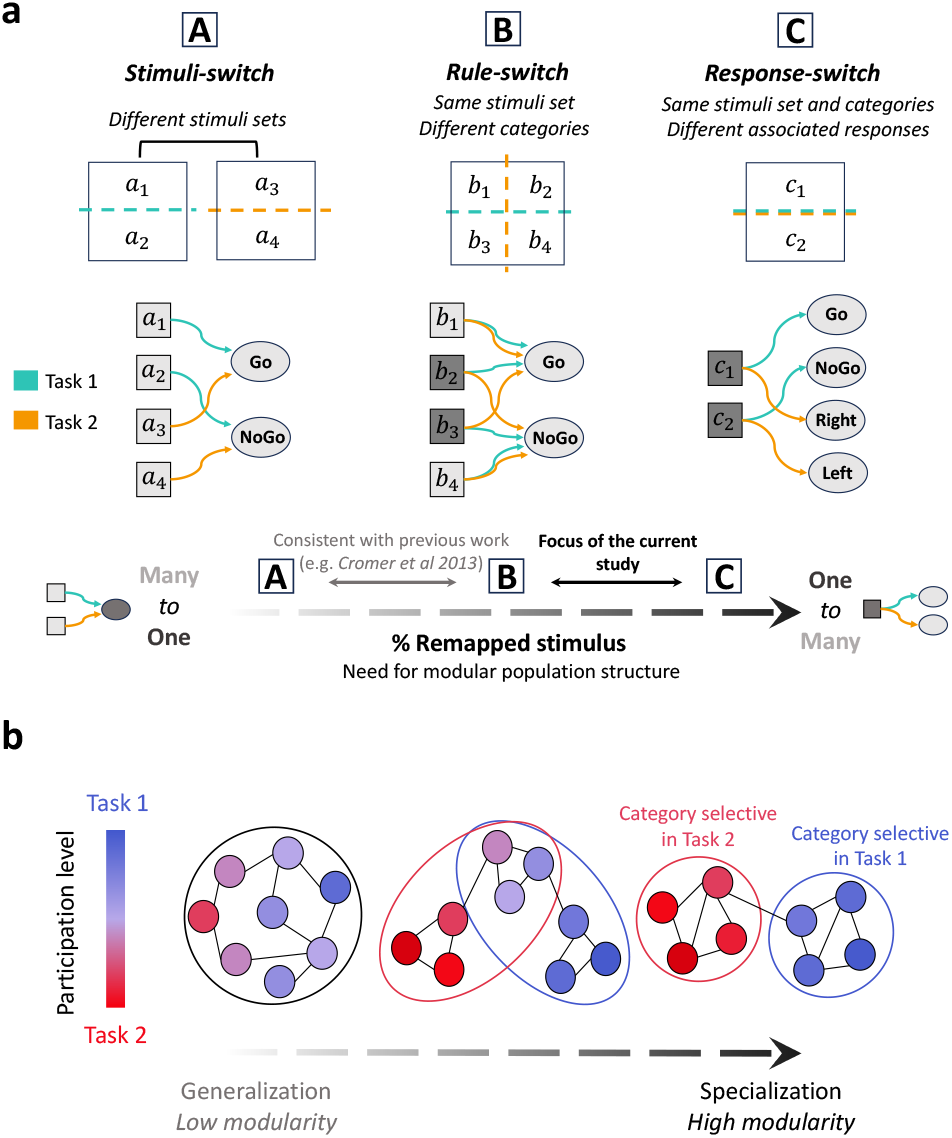
Increasing modularity of population structure with remapping requirements. **a**, Schematic of different context-dependent decision-making paradigms. Every paradigm is composed of two tasks. The first task is identical across all paradigms. An animal has to lick a spout (Go) in response to half of a stimulus set and refrain from licking (NoGo) for the other half. The second task, on the contrary, is different in each paradigm. (A) “Stimuli-switch”: in a second task with a stimulus set consisting of brand-new elements, the animal has to respond again Go to a subset and NoGo to the other. This requires mapping different inputs to the same output but no stimulus has to be remapped to different outputs depending on the task (0% remapping). (B) “Rule-switch”: as a second task the animal has to categorize the same set of stimuli using another rule. In this example, half of the stimuli has to be remapped to different output depending on the task (50% remapping). (C) “Response-switch”: as a second task the animal categorize the same set of stimuli with the same rule but has to map them to new outputs. In this example, the animal has to lick in a different direction depending on the stimulus category by either responding Left or Right. In this case, all stimuli have to be mapped to different output depending on the task (100% remapping). **b**, Schematic of population structure with different degrees of modularity. Left, only a large population of generalist neurons multiplexes representations of both tasks. This corresponds to a population structure composed of one module. Right, representations are split over two distinct populations of task-specialized neurons. This corresponds to a population structure with higher modularity. In this work, we hypothesize that the stripped axes from **a** and **b** are in fact aligned.

How does the neural code evolve from one task to another to handle these computational constraints? One possibility is that a generalist neural population is involved in representing inputs and outputs in both tasks. Another possibility is that representations are distributed across multiple task-specialized subpopulations, or modules, and that each task engages non-overlapping subsets of these modules [3] (Fig. 1b). Experimental data are compatible with both specialist and generalist neurons coding for task-relevant categories in prefrontal cortex (PFC), a cortical area known to play a critical role in cognitive flexibility [2]. While some studies support that different categorization tasks are mediated by different neural populations or circuits in PFC [4–8], others describe a substantial part of PFC neurons as multi-taskers that are involved in several categorization tasks [9–11]. Studies on recurrent neural networks (RNNs) have sought to address these apparent discrepancies. In particular, Dubreuil et al. (2022) [12] have suggested that a modular structure is required when identical stimuli have to be mapped to different outputs depending on the task. We therefore conjectured that the closer a paradigm is to the “one-to-many” end of the spectrum, the more prevalent task-specialized neurons should be, resulting in higher modularity for the population structure.

Nevertheless, most of the experimental studies have focused on many-to-one paradigms, leaving gaps in our understanding of the neural code underlying cognitive flexibility. In this study, we used a dual approach to tackle this question. First we tested whether modular structure can be found in a one-to-many paradigm. To this end, we analyzed calcium imaging data recorded in mouse mPFC [11] while the animals performed a visual categorization task with two different motor modalities (‘Response-switch’ paradigm, Fig. 1a, right). We discovered a modular population structure of three components: two subpopulations that specialized in a given task, and a third subpopulation that generalized across tasks. Secondly, and more importantly, there have been few investigations to date examining population structure across distinct task-switching paradigms. Here, by comparing data from a Response-switch and a Rule-switch paradigm (Fig. 1a, paradigms B and C), we aim to test whether modularity increases when performing tasks requiring more sensorimotor remapping. In line with our hypothesis, we observed less modularity in the Rule-switch paradigm compared to the Response-switch. Finally, from these experimental findings, we propose an input-driven mechanism based on gain modulation, which can explain how task-specialized neurons emerge to varying degrees depending on sensorimotor remapping requirements.

## 2 Results

### 2.1 Mouse mPFC activity during a Response-switch paradigm

To investigate mPFC population structure in flexible decision-making, we analyzed deconvolved calcium data previously collected in head-fixed mice (n = 9) that had to categorize visual stimuli into two groups [11] (Fig. 2a). Stimuli were 36 sinusoidal gratings, varying along two dimensions: orientation and spatial frequency. Either of these two features was used to define category boundaries for a given mouse (Fig. 2a, right). In each trial, a stimulus was presented for 1.5 seconds (Fig. 2b). The animals had to make a choice during this window. Mice were rewarded with water for correct answers, and received a time-out for incorrect ones.

**Fig. 2.**
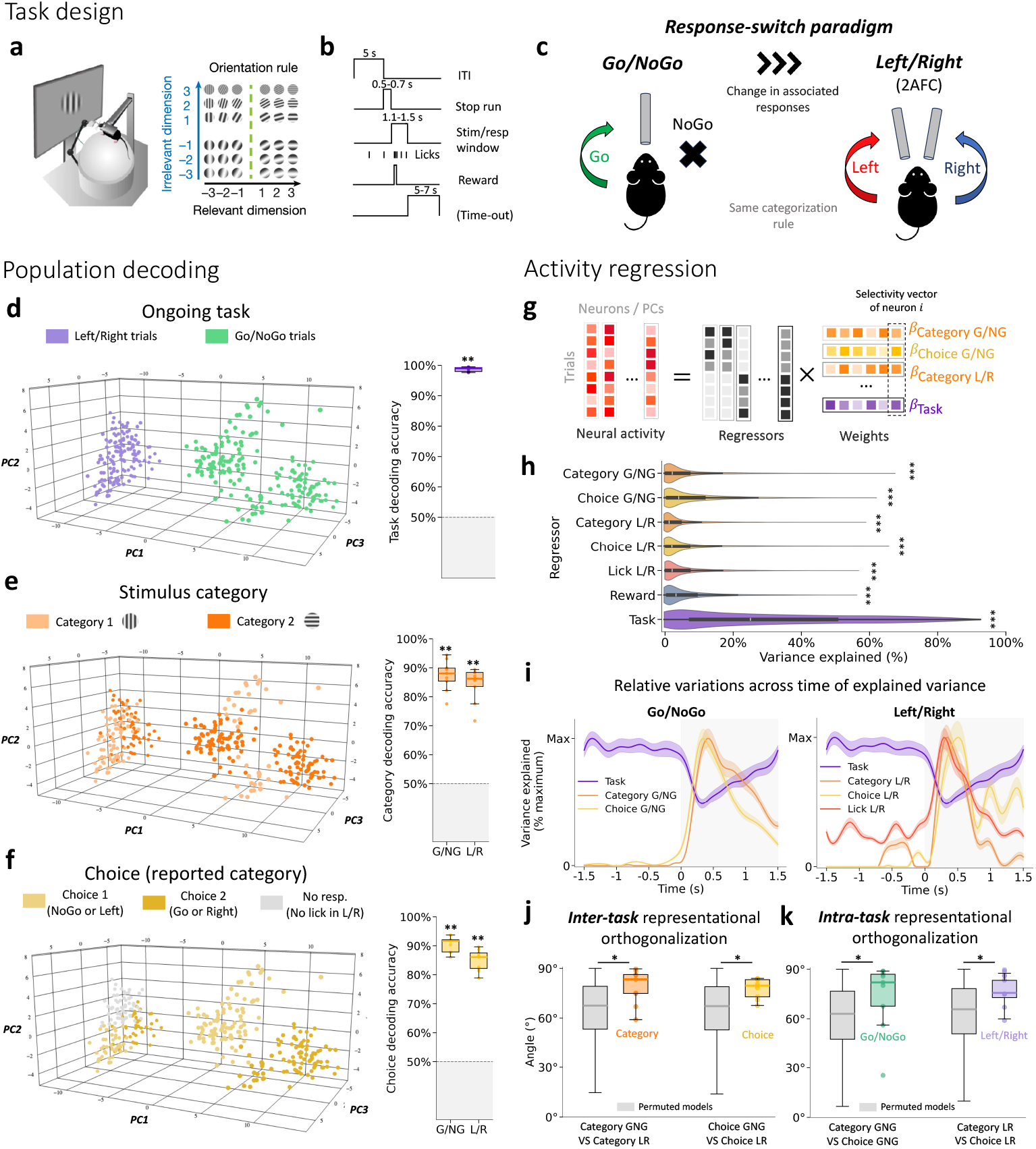
Geometry of mPFC representations at population level in a Response-switch paradigm. **a**, Left, schematic of behavioral training setup. Right, task design for an example mouse that had to categorize sinusoidal gratings based on their orientation. **b**, Schematic of trial structure. ITI, inter-trial interval; Stim/resp, stimulus presentation/response window. **c**, Schematic of the dual task paradigm. In a first task, mice had to either Go (lick a spout) or NoGo (refrain from licking) for the two stimulus categories, while they responded Left or Right (i.e. lick a spout either left or right) to the same categories in a second task. **d**, Linear decoding of task identity based on population activity, using Support Vector Classifiers (SVCs). Left, pseudo-population vectors for every pseudo-trial projected on a subset of first principal components. Right, distribution of decoding performance across all mice. Each dot corresponds to decoding performance for one animal. Decoding performance above chance was assessed with a Wilcoxon right-tailed signed rank test (** *P* < 10^−3^). **e**, Same as **d** for decoding of stimulus category. **f**, Same as **d** for decoding of animal choice. **g**, Linear regression of neural activity across the two tasks using multiple behaviorally-relevant regressors (G/NG:Go/NoGo, L/R:Left/Right). **h**, Variance explained by each regressor across all recorded neurons (corrected for chance). All regressors explained a significant amount of variance (*** *P* < 10^−4^, Wilcoxon right-tailed signed rank test, n=2389 neurons, Bonferroni corrected for seven comparisons). **i**, Variation over time of the mean explained variance across neurons by different regressors in the Go/NoGo task (left) and Left/Right task (right). To better visualize relative changes, variance explained is normalized to its maximum value for each regressor. Grey shaded area corresponds to the stimulus presentation window. Shaded areas around the curves correspond to 95% bootstrap confidence intervals. **j**, Left, orthogonalization of category and choice axes across animals when switching task. Orange, angles between the category axis in Go/NoGo and the category axis in Left/Right. Yellow, same for the choice axes. Each dot corresponds to one animal. Grey, distribution of angles between the same set of regressors for permuted models. **k**, Same as **j** for angles between category and choice axes computed in Go/NoGo (light green) or in Left/Right (light purple). Angle distributions for true regression axes were compared with angles from permuted models using a one-sided Mann-Whitney U test, Bonferroni-corrected for four comparisons (* *P* < 0.05).

Mice were initially trained in a Go/NoGo task to lick a spout for half of the stimuli (Go) and to refrain from licking for the other half (NoGo). Next, the same mice performed a Left/Right task, licking to the left for one category or to the right for the other category (Fig. 2c). Thus, while the stimulus set and category boundaries remained the same from one task to the next, their associated responses changed. We referred to such a design as a “Response-switch” paradigm (Fig. 1a, paradigm C), in which all stimuli have to be remapped between tasks. Identical neurons across the two tasks were recorded in mPFC with two-photon calcium imaging, allowing us to investigate how neural representations evolved during response switch, from Go/NoGo to Left/Right.

### 2.2 Population-level representations of task variables are non-interfering and substantially task-modulated

We first described how task variables were represented at the population level. To do so, we identified prevalent neural modes by performing dimensionality reduction of population activity with principal component analysis (PCA) (Fig. 2d). Despite the large size of the initial space (*n* = 2389 neurons across 9 mice), neural representations were embedded in a low-dimensional subspace (*d* ≤9, see Supplementary Fig. 1). We found that ongoing task identity (Fig. 2d), stimulus category (Fig. 2e) and animal choice (Fig. 2f) were highly decodable in both tasks across all individuals (*P*_*GNG*_ = *P*_*LR*_ = 0.0019, Wilcoxon right-tailed signed rank test, *n* = 9 mice). Interestingly though, we were unable to decode above chance stimuli belonging to the same category (*P* = 0.71, Wilcoxon right-tailed signed rank test, *n* = 9 mice, Supplementary Fig. 2), suggesting that representations of stimulus features were dominated by behaviorally relevant categories, consistent with studies proposing that PFC mainly receives early gated inputs [13–16].

We first set out to disentangle the contributions of stimulus category and animal choice into population-level representations. Indeed, given the animals’ good performance on the tasks (mean *d*′ ≥ 1.5) [11], stimulus category and animal choice were moderately correlated. To separate the contribution of choice and category in neural representations, we used ridge linear regression on the activity averaged over stimulus window (Fig. 2g). The resulting regression weights define encoding axes along which population activity carries information about task variables, enabling us to extract neural axes encoding for category, choice, licks, and task identity. Each regressor explained a significant part of mPFC activity variance (Fig. 2h). Task identity accounted for the most variance (30.6 ± 0.5%, *s*.*e*, n=2389 neurons) while other regressors explained smaller amounts on average (5.8 ± 0.2%, *s*.*e*). Explained variance was computed using cross-validated *R*^2^ (cv*R*^2^) score [17]. cv*R*^2^ accounts only for the variance that a regressor reliably captures, correcting for what would be expected by chance with a model of same size [17]. The variance explained by task identity and task variables evolved differently over the course of a trial. Category- and choice-related activities were stimulus-evoked, peaking between 300ms and 500ms after stimulus onset (Fig. 2i). In contrast, task-related activity was present before stimulus onset and remained relatively constant throughout a trial (Fig. 2i). A complementary cross-correlation analysis revealed that the neural axis encoding task identity remained stable over the course of a trial (Supplementary Fig. 3), while category and choice neural axes emerged only after stimulus presentation and were comparatively more time-varying.

Next, we investigated changes in task variable selectivity upon response switch. Neural representations of category and choice were strongly modulated between tasks, despite identical stimulus categories, with both translation (Supplementary Fig. 3a) and rotation of the associated neural axes (Fig. 2j,k, Supplementary Fig. 3a). More specifically, category and choice encoding axes were more orthogonal than expected by chance, both between (*P*_*Category*_ = 0.0044, *P*_*Choice*_ = 0.0089, one-sided Mann-Whitney U test, *n* = 9 mice, Fig. 2j) and within tasks (*P*_*G*/*NG*_ = *P*_*L*/*R*_ = 0.011, Fig 2k). This result supports the idea that mPFC encodes task variables across decorrelated channels to minimize interferences both between and within tasks [18–20]. While representational orthogonalization of categories in PFC have been reported in human fMRI and monkey single unit data [5, 15], our data suggest that this mechanism also extends to the representation of the ongoing choice.

### 2.3 mPFC neurons are most involved in the representation of variables in their preferred task

Even though the decorrelation of representations is informative about the way mPFC operates when dealing with conflicting tasks, this could occur without requiring a modular population structure. Indeed, the same neuronal population could achieve decorrelated representations across tasks by leveraging distinct patterns of activity [15, 19]. To determine whether task-dependent modulations of category and choice axes were supported by distinct subpopulations, we analyzed neural selectivity to task variables based on regression weights. We found that the regression weight for the task, which we term *task selectivity*, polarized the participation of neurons between tasks.

Time variation of neural activity over the course of a trial was highly task-specific and structured by task selectivity. In their preferred task, neurons’ activity varied more over time (Fig. 3a). Although these neurons were more involved in temporal dynamics, we wanted to verify that these variations reflected the representation of variables useful to the task, rather than noise. As a result, we inspected category-dependent changes in peri-stimulus histograms for each neuron. Task-preferring neurons (|*β*_*Task*_| > 0) showed larger category-dependent changes in post-stimulus activity in their preferred task (Fig. 3b). To quantify the overall participation of neurons in task-relevant representations, we introduced a *task participation* score, averaging the absolute regression weights for category and choice within each task (see Supplementary Fig. 4a for details). Again, we found that inter-task variability (captured by task selectivity) translated into greater intra-task variability (captured by task participation): neurons specialized by predominantly representing category and choice in their preferred task (Fig. 3c). Our observation is consistent with previous studies. Theoretical work on RNNs predicted a correlation between selectivity for the task and selectivity for stimulus features [12]. Experimental studies further support that task-dependent increase in pre-stimulus firing rate correlates with increase in evoked firing rate in both primary auditory cortex and PFC [21]. Additionally, we observed a symmetric decrease in task participation at both extremity points for highly task preferring neurons (|*β*_*Task*_| *≈* 1). This effect could not be explained by the regularization involved in ridge regression alone since it appeared even for models trained without regularization (Supplementary Fig. 4b). We propose in section 2.6 that this effect is due to the saturation of neurons by the task signal.

**Fig. 3.**
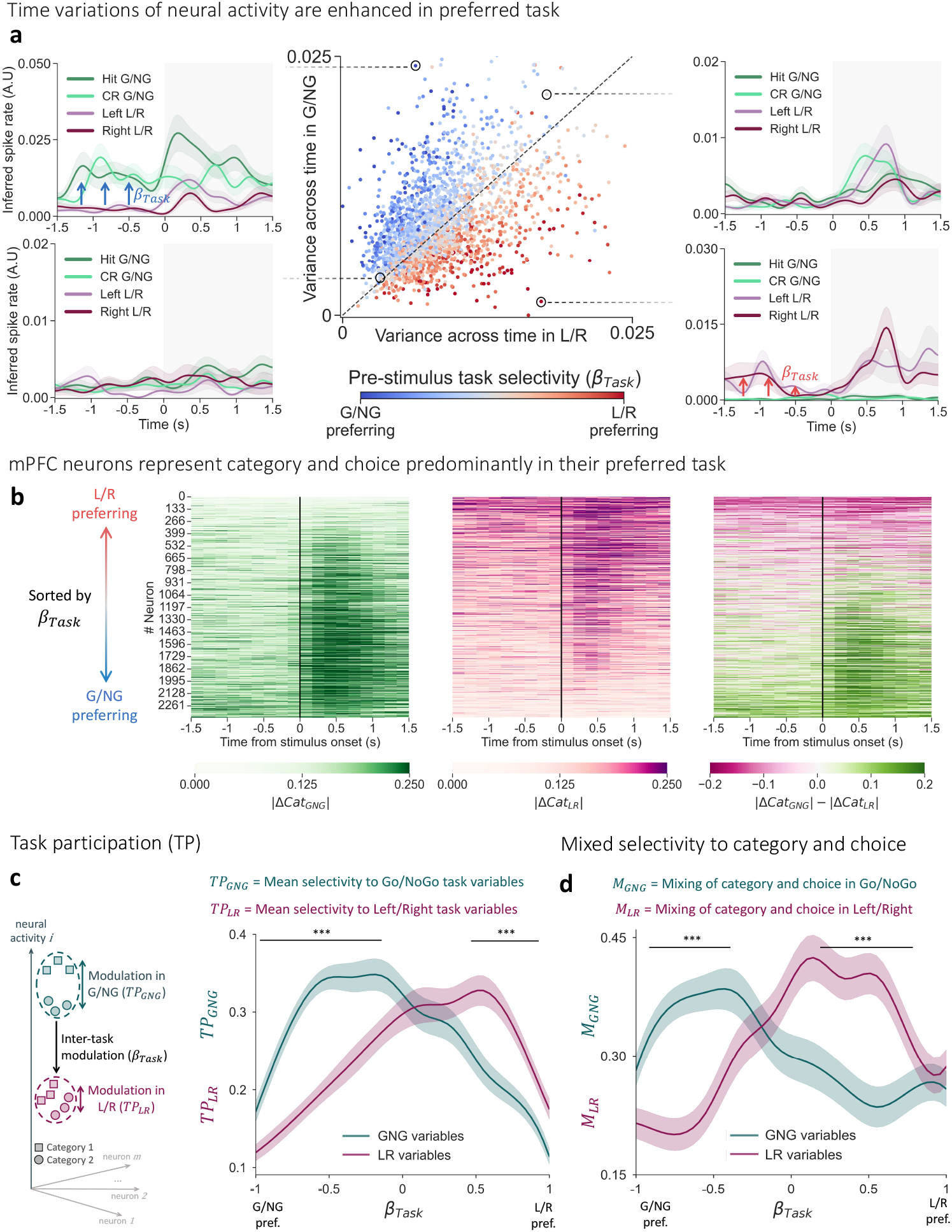
Representation of task variables by task-preferring mPFC neurons. **a**, Center, activity variance of individual neurons over time in Go/NoGo trials vs. Left/Right trials. Regression weight for the task (*β*_*Task*_) of corresponding neurons is represented by dot’s color. Dotted line corresponds to *y* = *x* axis. Left and right, Peri-stimulus time histograms of four example neurons. Grey shaded area indicates stimulus presentation window. **b**, Heatmaps highlighting category-dependent activity differences (Z-score). Neurons (rows) are sorted according to their task selectivity. Left, mean absolute activity difference in Go/NoGo between trials with a presented stimulus from category 1 versus category 2. Center, same for trials from Left/Right task. Right, left heatmap minus center one highlighting a task-dependent sensitivity to the stimulus category for highly task preferring neurons. **c**, Task participation scores for each task as a function of task selectivity. Participation scores were computed as the mean absolute selectivity for task variables (Category + Choice) from Go/NoGo (dark green) or from Left/Right (magenta). **d**, Cosine similarity between selectivity vectors and the *y* = *x* axis. Neurons aligned with *y* = *x* axis (high cosine similarity) are selective in the same proportion to both stimulus category and animal choice, situation we refer as high mixing of category and choice. Mixing of category and choice in Go/NoGo (dark green) and in Left/Right (magenta) are plotted as a function of task selectivity (as in **b**). Shaded areas around the curves correspond to 95% confidence intervals (± 2 *s*.*e*, standard error). Differences between the two curves in **b** and **c** where assessed using a two-sided Mann-Whitney U tests, Bonferroni-corrected for multiple comparisons (*** *P* < 10^−4^).

Next, we aimed to assess the relative balance between stimulus category selectivity and choice selectivity across neurons by examining their mixed selectivity, a common feature in PFC [22, 23]. Interestingly, task-selective neurons exhibited strong mixed selectivity for category and choice in their preferred task (Fig. 3d). This suggests that task-specialized neurons play a critical role in integrating category and choice in a task-specific manner [24], a key step to flexibly map stimuli to appropriate motor output. To ensure that these results were not due to unanticipated circularity caused by the regression, our analyses were cross-validated with another model trained on separate time periods. In that model, task selectivity was obtained by performing regression on neural activity averaged over a 1.5s pre-stimulus window, while selectivities to other task variables, such as category and choice, were derived from post-stimulus activity (1.5s). Both task participation and category-choice mixing effects remained present under this control (Supplementary Fig. 4c,d).

### 2.4 Task selectivity delineates three neural subpopulations with distinct functional roles

We next investigated whether the strong structuring effects of task selectivity resulted in functionally distinct neural subpopulations. To this end, we assigned each neuron a selectivity vector consisting of its regression weights in the two tasks, representing a neuron’s selectivity as a point in a space with seven dimensions corresponding to the number of regressors. We then asked whether these selectivity vectors were randomly distributed or whether they clustered in structured modules.

We identified three subpopulations in mPFC (Fig. 4a), primarily structured by task selectivity (Fig. 4b). Two out of three subpopulations were highly task preferring: one preferred Go/NoGo (Population GNG) and the other Left/Right (population LR) (*P* < 10^−20^, two-sided Mann-Whitney U test, *n*_*GNG*_ = 509, *n*_*LR*_ = 405 neurons). The third subpopulation (Population 0) lacked a consistent task preference and spanned a broader range of selectivity. These subpopulations were identified by fitting a Gaussian Mixture (GM) model to the 7-dimensional point cloud. GM models seek to describe an observed distribution by combining multivariate Gaussian distributions in an unsupervised manner. These models have the advantage of being highly interpretable, as they provide components that can be directly identified as subpopulations of the initial distribution. A three-component model provided the best fit to the data, as adding more clusters did not significantly improve likelihood (Fig. 4c). To further quantify local structures in selectivity space we applied an ePAIRS test, a non-parametric method that evaluates the angular distribution of vectors in the point cloud against a null distribution. ePAIRS revealed that selectivity vectors were significantly more clustered than expected by chance (*P* = 5.0 × 10^−5^, Fig. 4d), indicating that the distribution of our data had multiple preferred directions, incompatible with a Gaussian random structure. Consistent with this, a large effect size underscored a high degree of modularity in the selectivity space (Fig. 4e).

**Fig. 4.**
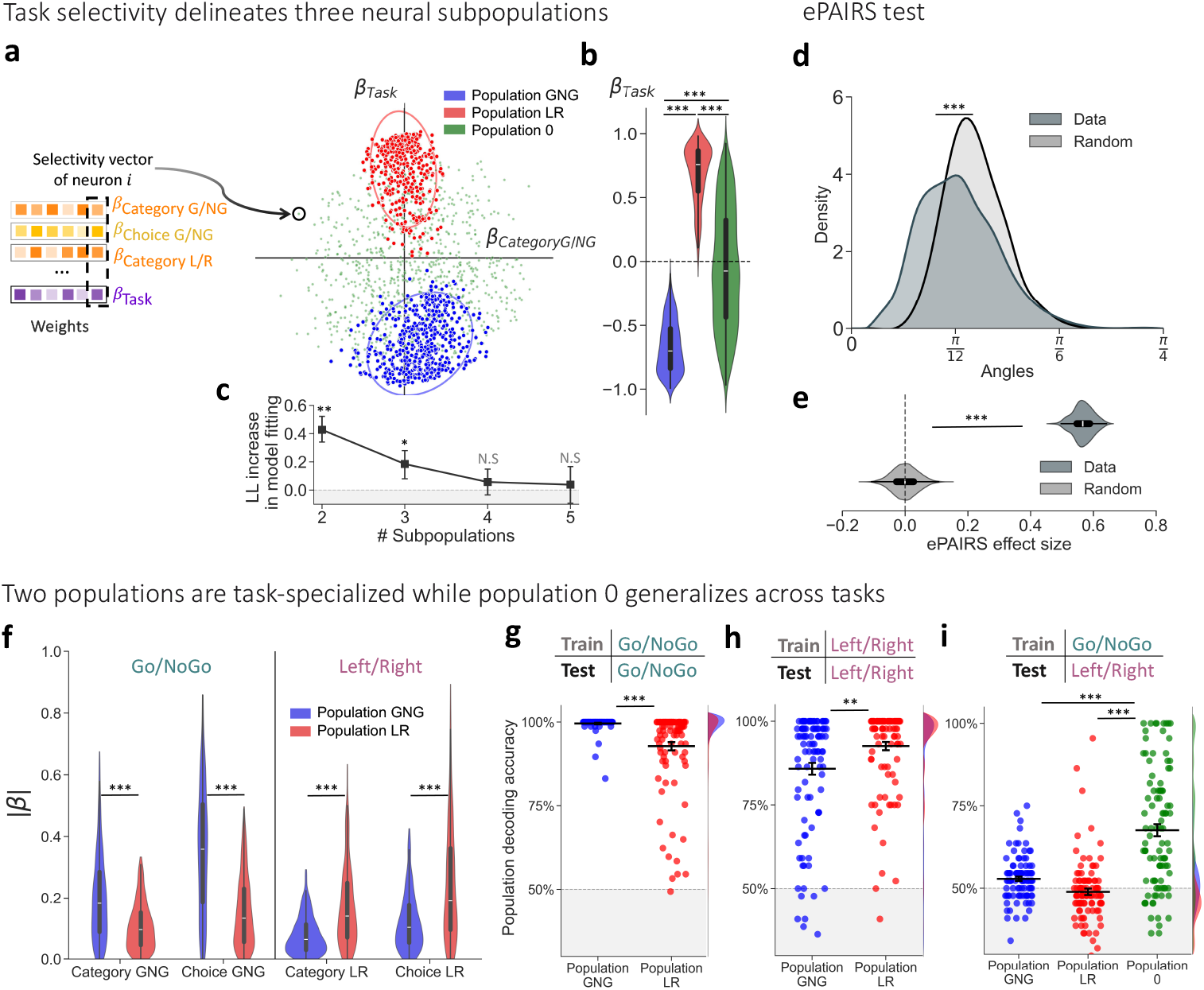
Clustering of neural subpopulations by selectivity to task variables. **a** 2-D projections of individual selectivity vectors for a combination of two example regressors (Category in Go/NoGo and Task). Colors indicate which population extracted by the Gaussian mixture (GM) model a neuron belonged to (see Methods). Ellipses indicate 95% confidence intervals. **b**, Task selectivities (*β*_*Task*_) for neurons from the three identified populations. Population GNG was Go/NoGo preferring (*β*_*Task*_ < 0) while Population LR was Left/Right preferring (*β*_*Task*_ > 0). Differences in task selectivity between the populations were assessed with a two-sided Mann-Whitney U test, Bonferroni-corrected for three comparisons (***, *P* < 10^−4^). **c**, Increase of the per-sample average log-likelihood with each component added to the model. Grey shaded area corresponds to a lack of improvement in model performance. Log-likelihood stopped increasing significantly after the addition of a third component. Significant increase in model performance was assessed using a one-sided permutation test Bonferroni-corrected for multiple comparisons (N.S : Not significant, * *P* < 0.05, ** *P* < 0.001). **d**, Distribution of angles between each selectivity vector and its nearest neighbor (Dark blue, ‘Data’) compared to a null-distribution constructed from Gaussian random vectors with the same statistics as the data (Grey, ‘Random’). The mismatch between the two distributions was assessed using the ePAIRS test (one-sided, see Methods). **e**, Effect sizes of the ePAIRS test for different null-distributions. The effect size of ePAIRS test in the data was greater than expected by chance (*P* < 10^−4^, two-sided Mann-Whitney U test). **f**, Absolute selectivity to task variables from Go/NoGo and from Left/Right for neurons from Population GNG (blue) or from Population LR (red). Differences in absolute selectivity between the populations were assessed for each regressor with a two-sided Mann-Whitney U test, Bonferroni-corrected for five comparisons. **g**, Trial-based linear decoding of the category of the stimuli on correct trials with neurons from different populations. Decoders were Support Vector Classifiers (SVCs) trained on half of the trials from Go/NoGo and tested on the other half. Classifier used non-overlapping subsets of 2 neurons drawn without replacement from Population GNG (blue) or from Population LR (red) **h**, same as in **g** for classifiers trained and tested on Left/Right trials. **i**, same as in **g**-**h** for classifiers trained on half of the trials from Go/NoGo and tested on half of the trials from Left/Right (for comparable train/test set size with **g**-**h**). Classifiers used non-overlapping subsets of 20 neurons from a given population. Differences in decoding accuracy between the populations were assessed with a two-sided Mann-Whitney U test, Bonferroni-corrected for four comparisons.

Each subpopulation had distinct functional roles, explaining the observed interaction between task preference and task participation. Task-specialist populations GNG and LR were primarily involved in encoding task-relevant information for their preferred task with greater selectivity (Fig. 4f). The functional specialization of these populations was further supported by decoding analyses using support vector classifiers (SVCs). Stimulus category was decoded more accurately when classifiers were trained using the specialist population (*P*_*GNG*_ = 5.1 × 10^−13^, *P*_*LR*_ = 1.6 × 10^−4^, two-sided Mann-Whitney U test, *n* = 100 decoders, Fig. 4g,h). We further examined how representations of stimulus category generalized across tasks. Since mice were trained chronologically on Go/NoGo and then on Left/Right, we compared decoding performance when training on Go/NoGo and testing on Left/Right trials. Only Population 0 exhibited significant generalization across tasks (*P* = 6.3 × 10^−13^, Wilcoxon right-tailed signed rank test), implying a shared neuronal code across tasks (Fig. 4i). These results suggest that Population 0 supports a generalist, i.e. task-invariant, encoding of stimulus category, providing robust and transferable neural representations across behavioral contexts. In contrast, specialist populations exhibit task-dependent category encoding, facilitating context-specific processing of categorical information.

### 2.5 Modularity of mPFC population decreases in a paradigm with less sensorimotor remapping

Altogether, the above results support the hypothesis that mPFC population structure is organized into task-specialized modules to perform tasks that involve conflicting associations. Next, we investigated whether a fixed level of modularity is sufficient to handle varying remapping requirements in an all-or-none manner, or whether modularity scales with the number of stimuli to be remapped. To address this, we analyzed a separate dataset where a different cohort of mice (n = 12) was trained on another task paradigm using the same stimuli. As before, mice were first trained on a Go/NoGo task to categorize sinusoidal gratings by orientation or spatial frequency. They were then retrained on a second task to categorize the stimuli by the alternate feature while maintaining the Go/NoGo response (Fig. 5a). We refer to such a design as a “Rule-switch” paradigm (Fig. 1a, paradigm B). In this case, only half of the sensorimotor associations conflicted between tasks, resulting in less sensorimotor remapping compared to the Response-switch paradigm.

**Fig. 5.**
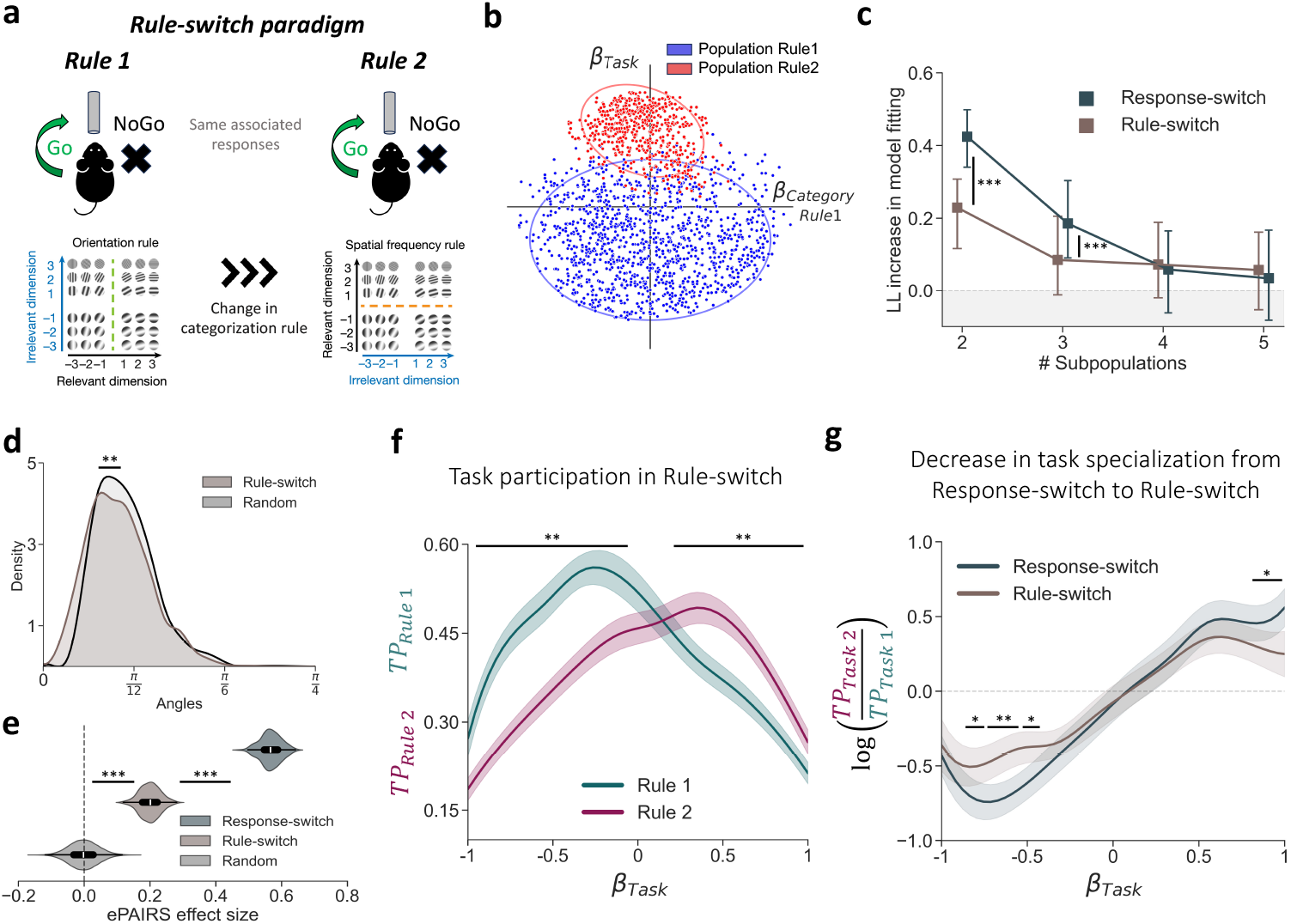
Comparison of population modularity in Response-switch and Rule-switch paradigms. **a**, Schematic of a Rule-switch dual task paradigm (Fig. 1a, Paradigm B). In both tasks, mice categorized sinusoidal gratings with a Go/NoGo design. In a first task (Rule 1), they categorized gratings by orientation, while in a second task (Rule 2), they categorized gratings by spatial frequency. **b**, 2-D projections of individual selectivity vectors for a combination of two example regressors (Category in Rule 1, and Task). Colors indicate which population extracted by the Gaussian mixture (GM) model a neuron belonged to (see Methods). Ellipses indicate 95% confidence intervals. **c**, Increase of the per-sample average log-likelihood with each component added to the model. Results from Rule-switch (Brown) are compared to those from Response-switch (Fig. 4e) represented in dark blue. Grey shaded area corresponds to a lack of improvement in model performance. Log-likelihood stopped increasing significantly after two components for Rule-switch compared to three for Response-switch (one-sided permutation test, see Methods). Differences in log-likelihood increase by each additional component were compared between Rule-switch and Response-switch with a two-sided Mann-Whitney U test (*** *P* < 10^−4^). **d**, Distribution of angles between each selectivity vector for Rule-switch and its nearest neighbor (Brown, ‘Data’) compared to a null-distribution constructed from Gaussian random vectors with the same statistics as the data (Grey, ‘Random’). The mismatch between the two distributions was assessed using the ePAIRS test (one-sided, see Methods). **e**, Effect sizes of the ePAIRS test for different null-distributions. The effect size of ePAIRS test in Rule-switch was greater than expected by chance but still significantly smaller compared to Response-switch (two-sided Mann-Whitney U test). **f**, Mean absolute regression weights for task variables from Rule 1 (Dark green) and from Rule 2 (Magenta), plotted as a function of the regression weight for the task (i.e the rule). We referred to these curves as ‘participation scores’ (Dark green: Participation in Task 1, Magenta: Participation in Task 2). **g**, Comparison of task specialization of neurons from Response-switch (dark blue) with neurons from Rule-switch (brown). Task specialization was computed as the log-ratio between participation scores for each task in a given paradigm and are plotted as a function of task selectivity (*β*_*Task*_). Neurons that participate in the same proportions in both tasks have a specialization close to 0, while neurons that participate preferentially in one task have a specialization big in absolute value (< 0 specialized in first task, > 0 specialized in second task). Shaded areas around the curves correspond to 95% confidence intervals (± 2 *s*.*e*). Differences in task participation (from **f**) and task specialization (from **g**) where assessed with a two-sided Mann-Whitney U tests, Bonferroni-corrected for multiple comparisons (* *P* < 0.05, ** *P* < 0.001).

mPFC population structure in Rule-switch exhibited less modularity and was best described by only two subpopulations (Fig. 5b,c). However, as in Response-switch, task selectivity remained the main factor delineating these subpopulations (Fig. 5b). ePAIRS analyses confirmed that selectivity vectors in Rule-switch were more randomly distributed than in Response-switch, although still deviating from a null distribution (*P* = 6.8 × 10^−8^, one-sided Mann-Whitney U test, Fig. 5d,e). In addition, we observed again a strong interaction between task preference and task participation indicating that the two identified subpopulations were task-specialized (Fig. 5f). However, neurons from both subpopulations were significantly less specialized than for Response-switch (Fig. 5g), suggesting that modularity scales with remapping requirements.

### 2.6 Population modularity can be explained by an interaction between task signal and global inhibition

Despite quantitative differences, we consistently observed task-specialized neurons across both Response-switch and Rule-switch paradigms. This suggests that task specialization may not strictly result from connectivity patterns learned through a specific set of tasks. We hypothesized instead that task specialization emerged from input-driven mechanisms, specifically the integration of stimulus and task signals by network units. To test this, we modelled several recurrent neural networks with randomly generated connectivity and asked if task specialization could be described across distinct recurrent structures only by fitting input statistics. Moreover, we accounted for the extensive body of evidence that context-dependent processing in biological networks is not only determined by external inputs but is also modulated by internal states that globally influence neural excitability. In sensory cortices, such modulation —often linked to attention, arousal, or task engagement— adjusts response gain by enhancing responses for target inputs and suppressing irrelevant ones [25–28]. Interestingly though, such state-dependent modulations are not limited to early sensory cortices and may intensify along the cortical hierarchy, peaking in higher-order areas [25, 29, 30]. We therefore tested whether state-dependent activity modulation could explain task specialization in our mPFC model by introducing a *global modulation* input that sets neuron baseline activity upon engagement in a paradigm (Fig. 6a). This global modulation acts as a bias, effectively shifting neurons’ activation functions leftward or rightwards, depending on its sign. Global modulation differs from the ongoing task input both spatially and temporally. Spatially, the task signal is heterogeneously fed into the network with balanced excitatory and inhibitory weights centered around zero, whereas global modulation has non-zero mean projections weights, leading to an overall excitatory or inhibitory effect across the network. Temporally, the task signal flips in sign between tasks, whereas global modulation remains constant across the two tasks of a paradigm.

**Fig. 6.**
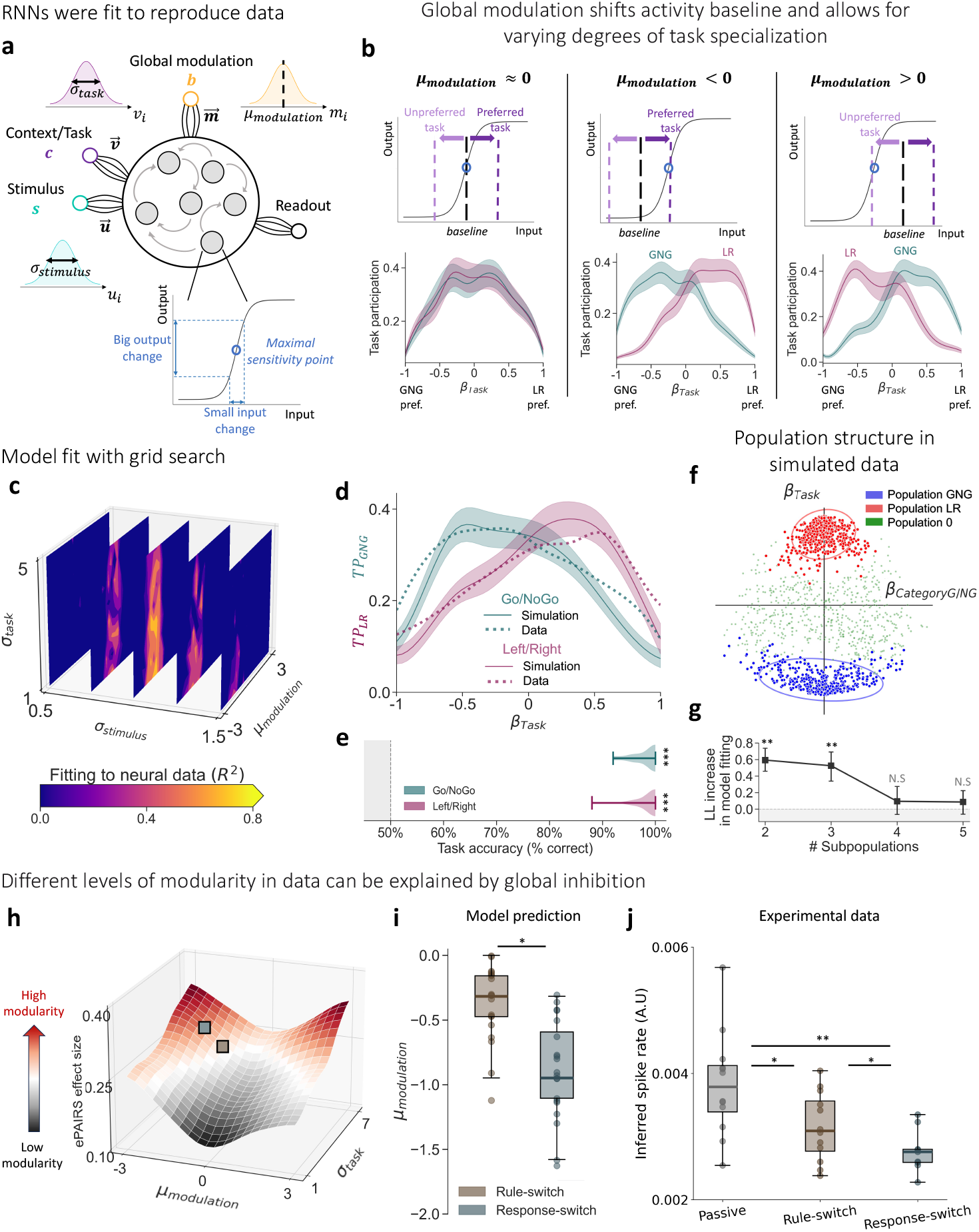
A model of task-specialized neural modularity driven by global modulation and task signals. **a**, Recurrent neural networks with random connectivity received noisy inputs about task identity (purple) and stimulus category (cyan) in addition of a global modulation input (orange) homogeneously fed to each unit. The model was fitted to reproduce the task participation curves found in the data by varying the standard deviation of projections weights for task signal (*σ*_*task*_) and stimulus signal (*σ*_*stimulus*_) and the mean projection weight for global modulation (*µ*_*modulation*_). Activation function of model units was a sigmoid function. Sensitivity of a unit’s output to its input (or gain) changes as a function of the input with a maximum at inflection point (represented with a blue circle). **b**, Task participation of model units for different values of *µ*_*modulation*_. Left, without global modulation neural gain decreased symmetrically in both tasks leading to no specialization. Center, by globally inhibiting neurons their baseline is pushed away from maximum sensitivity regime. Task signal restores neural sensitivity in a task-dependent manner leading neurons to better represent task variables in their preferred task as observed in the data (Fig. 3c). Right, A global excitation led to the opposite effect with neurons specializing in their unpreferred task. However, we keep the decline in task participation for highly task-preferring neurons (|*β*_*Task*_| *≈* 1) that was observed in data. This is due to task signal pushing neurons to low sensitivity regime. **c**, Heatmap of model fitting performance (computed as *R*^2^ score) on Response-switch data for different (*σ*_*stimulus*_, *σ*_*task*_, *µ*_*modulation*_) combinations. Best fit is framed in black. **d**, Participation curve for each task in Response-switch with neurons from the best-fit model (solid line) compared to neurons from data (dotted line). Shaded areas around the curves corresponds to ± 2 *S E*. **e**, Performance on the two tasks from Response-switch by training a linear readout on the network that best fits task participation curves. **f**, 2-D projections of individual selectivity vectors, from model units with best fit to Response-switch data, for a combination of two example regressors (Category in Go/NoGo, and Task). Colors indicate which population extracted by the Gaussian mixture (GM) model a neuron belonged to. Ellipses indicate 95% confidence intervals. **g**, Increase of the per-sample average log-likelihood with each component added to the GM model. Grey shaded area corresponds to a lack of improvement in model performance. Log-likelihood stopped increasing significantly after the addition of a third component as observed in the data (Fig. 4c). Significant increase in model performance was assessed using a one-sided permutation test Bonferroni-corrected for multiple comparisons (N.S : Not significant, ** *P* < 0.001). **h**, ePAIRS effect size for different (*σ*_*task*_, *µ*_*modulation*_) combinations. Red regions correspond to models composed of units with highly clustered selectivity vectors. **i**, *µ*_*modulation*_ that provides the best fit to data in Rule-switch (brown) or in Response-switch (dark blue) for 20 different iterations of model connectivity. Best *µ*_*modulation*_ was significantly more negative in Response-switch compared to Rule-switch (*, *P* < 0.05, two-sided Mann-Whitney U test). **j**, Mean inferred spike rate across all recorded neurons for different conditions. Each individual is represented by a dot. In “passive” sessions mice were not yet engaged in the task but were presented with the same stimulus set as in Rule-switch and Response-switch sessions. For Response-switch and Rule-switch neural activity was averaged across sessions from both tasks composing a paradigm.

Each unit was modeled with a non-linear sigmoid activation function with maximal sensitivity (i.e. derivative) at zero (Fig. 6a). By shifting the input baseline away from maximal sensitivity point, global modulation induced asymmetric sensitivity across tasks. This asymmetry proved crucial in explaining both the direction and magnitude of the interaction between task selectivity and task participation in our data. Negative modulation (*µ*_*modulation*_ < 0, Fig. 6b) led neurons to specialize in their preferred task, aligning with the data. In contrast, positive modulation (*µ*_*modulation*_ > 0, Fig. 6b) led to specialization in the unpreferred task. Meanwhile, task specialization disappeared in the absence of modulation (*µ*_*modulation*_ *≈* 0, Fig. 6b). For moderate modulation values (|*µ*_*modulation*_| < 3), we observed a symmetric decrease in task participation for highly task preferring neurons (|*β*_*Task*_| *≈* 1), consistent with the data (Fig. 3c). This pattern suggests that saturation of neural activity by strong task inputs could underlie the decrease in participation observed at the extreme points of the curves.

A network integrating stimulus, task and global modulation inputs was able to replicate the main findings of our analysis on both Response-switch (Fig. 3 and 4) and Rule-switch (Fig. 5) paradigms. We fitted the model to reproduce task participation curves (Fig. 3c) using a grid search procedure over three parameters: stimulus and task signal strength (*σ*_*stimulus*_, *σ*_*task*_) and mean global modulation across the population (*µ*_*modulation*_). Adjusting these parameters allowed the model to closely fit the data (*R*^2^ = 0.76) (Fig. 6c,d) and perform the two tasks in a paradigm (Fig. 6e). Both *σ*_*stimulus*_ and *µ*_*modulation*_ were critical in the fitting process, with good scores only within a narrow range of values (Fig. 6c). In contrast, the model achieved good scores for a wider range of *σ*_*task*_, provided the task signal was strong enough to drive task selectivity. Gaussian mixture analysis on the best-fit model for the Response-switch paradigm revealed three subpopulations delineated by task selectivity, as observed in the data (Fig. 6f,g).

While *σ*_*task*_ was not critical for reproducing the interaction between task selectivity and task participation, it played a crucial role alongside with *µ*_*modulation*_ to generate different degrees of modularity in population structure. Specifically, ePAIRS effect size increased with the product of *σ*_*task*_ and *µ*_*modulation*_ (Fig. 6h), indicating that modularity emerged from their interaction. In line with this result, we found that *µ*_*modulation*_ was critical for explaining the different degrees of modularity observed across paradigms. Networks fitted to task participation in Response-switch required stronger inhibition than those fitted to Rule-switch data (*P* = 5.6 × 10^−3^, two-sided Mann-Whitney U test, *n* = 20 networks, Fig. 6i). To evaluate whether the inhibitory modulation inferred by the model was supported by empirical data, we compared average neural activity across “passive”, “Rule-switch”, and “Response-switch” sessions. In “passive” sessions, mice were exposed to the stimulus set for the first time and were not yet engaged in the task, as indicated by minimal behavioral responses (no licking observed over 70% of the trials). We observed a systematic decrease in average activity from passive to task-engaged conditions (*P*_*Rule*−*sw*._ = 0.013, *P*_*Resp*−*sw*._ = 0.0016, one-sided Mann-Whitney U test, *n*_*Passive*_ = *n*_*Rule*−*sw*._ = 12, *n*_*Resp*−*sw*._ = 9 mice, Fig. 6j), consistent with our initial hypothesis that task engagement modulates activity in mPFC. Notably, this decrease was most pronounced during Response-switch sessions compared to Rule-switch (*P* = 0.047, one-sided Mann-Whitney U test, *n*_*Rule*−*sw*._ = 12, *n*_*Resp*−*sw*._ = 9 mice), in line with the stronger inhibitory modulation (i.e. lower *µ*_*modulation*_ values) inferred by the model for this condition. Taken together, these results suggest that global inhibition could be a key mechanism to control the degree of modularity in networks and promote the separation of conflicting representations into distinct subpopulations. This observation is biologically plausible, as it aligns with experimental evidence of cholinergic activation of cortical inhibitory interneurons during context-dependent behaviors, when comparing passive and task-engaged states [26].

## 3 Discussion

In this study, we investigated the impact of increasing sensorimotor remapping on the modularity of population structure in the mouse mPFC. By comparing two different paradigms —one with total remapping of inputs (Response-switch) and another with partial remapping (Rule-switch)—, we found that an increased need for remapping drives higher modularity in mPFC population structure. Through modelling, we proposed that a global modulation of population activity could act as a potential mechanism for regulating modular organization in a network depending on remapping requirements. Our results suggest that the modularity observed in our data could arise from the interaction between task-specific signals and global inhibition.

### Functional role of inhibition in cortical networks

In studies comparing task-engaged and passive states in trained animals[25–27, 31], task-specific signals are often confounded with broader attention- and arousal-related processes elicited by behavior. Here, by analyzing neural activity across two distinct tasks, we extracted task-specific activity and showed that it defines functionally distinct population modules. We further dissected this mechanism with a recurrent neural network that could reproduce modular structure in the population. Interestingly, we discovered that a global modulation shifting neurons’ baseline activity was sufficient to explain the emergence of modular population structure. This global modulation aligns with the differences in activity observed between task-engaged and passive states in previous studies [25–27, 31, 32]. Incorporating a global modulation in our model was motivated by growing evidence that internal states exert widespread influence on neural activity across cortical areas [33]. Such state-dependent modulation, which can be inhibitory or excitatory, often acts globally across neural population, resulting in substantial changes in average activity [25–28, 31, 32, 34–36].

Our experimental and modelling results predict that this global modulation should be predominantly inhibitory to explain the widely observed specialization of mPFC neurons in their preferred task. This conclusion is consistent with prior reports of reduced frontal cortical activity during task engagement [32] and heterogeneous recruitment of inhibitory interneurons across internal states in mice [26, 36]. However, it contrasts with findings in a related study where state-dependent modulation of sensitivity was mediated by an excitatory drive in the rat primary auditory cortex [16]. This discrepancy suggests that PFC may rely on inhibitory mechanisms to achieve functional modularity, highlighting a region-specific strategy for flexible neural computations. In any case, we showed that both inhibitory and excitatory modulation can drive modularity, provided they shift the neural input-output function toward a lower sensitivity regime, allowing the integration of stimulus category and task identity by network units.

Our proposed inhibitory mechanism is supported by the established role of mPFC as the principal hub of the default mode network (DMN), a system marked by elevated activity during rest and reliably suppressed during externally oriented, goal-directed behaviors. Recent advances in rodent neuroimaging have established a DMN analog in the mouse brain, wherein the mPFC, along with retrosplenial and lateral parietal cortices, exhibits synchronised low-frequency activity during rest that diminishes with task engagement [37, 38]. In both humans and mice, tasks requiring heightened attentional control or external sensory processing are associated with greater suppression of mPFC activity [39–42].

In the present study, we provided evidence that the deactivation of mPFC serves a functional role in shaping population structure during task engagement. Specifically, we showed that global inhibition promotes modularity in neural representations. This occurs via the dynamic segregation of distinct neural subpopulations, enabling them to function independently under the influence of task signal. We further linked variations in modularity across paradigms to different levels of global inhibition, suggesting that inhibitory networks can regulate the balance between generalization and task specialization, optimizing neural architecture to resolve task conflicts. Such a mechanism is both effective and biologically convenient, as it relies on coarse inhibitory drive rather than fine-tuned synaptic structures. By enabling flexible reconfiguration of population codes with minimal architectural constraints, this broad modulation offers a scalable and robust strategy for adaptive neural computation that may be readily implemented across diverse brain regions to support context-dependent processing.

### Comparisons to population structures found in previous studies

While multi-population structure have been found in artificial neural networks (ANNs) performing context-dependent tasks [12, 16], these studies relied on connectivity weights between units and could fully access the underlying network architecture, making direct comparisons with experimental neural activity challenging. Still, we sought to test predictions derived from connectivity-based ANN studies using neural activity alone. Clustering neurons according to their selectivities yielded conclusions that were largely consistent with connectivity-based ANN analyses. Interestingly though, we found multiple subpopulations in selectivity space despite our model consisting of a single neural population in connectivity space. This suggests that while subpopulations in connectivity space may translate into subpopulations in selectivity space, distinct subpopulations in selectivity space can be observed without structure in the connectivity space. Overall, our study supports an extension of connectivity-based population decomposition to activity-based analysis [43]. While such an extension holds great promise as a tool to identify functionally distinct neural ensembles, it currently lacks the theoretical foundations that underpins connectivity-based population decomposition [44]. In this regard, we believe that further analytical work exploring links between these two types of population decomposition would be highly valuable and would promote inter-disciplinary dialogues.

A similar three population structure has been found in the rat primary auditory cortex for a paradigm analogous to the Rule-switch condition [16], suggesting that task specialization could be a general principle expressed to varying degrees along the sensorimotor pathway. Why then do task-specialized subpopulations emerge in sensory areas under low-conflict conditions, and less so in prefrontal cortex under similar task demands? One possible explanation lies in the nature of the transformation performed in each region. In sensory cortices, task specialization may reflect the conversion of the stimulus set into abstract categories, depending on contextual demands. In contrast, the PFC may instead control the mapping of such abstract categories onto rule-appropriate actions, supporting behavioral flexibility across tasks. Our results suggest then potential roles for the different functional subpopulations identified in PFC. The generalist population (population 0), which is active across tasks, may serve as an intermediate layer that gathers categorical information, e.g., stimulus identity or decision variables, from upstream regions, and routes it towards task-specialized populations. These downstream units would then transform the abstract category into a task-specific motor action. This organization could reflect a hierarchical division of information processing optimized for context-dependent behaviour.

### Conflicting tasks and advantages of modular population structure

When outlining our framework, we defined the degree of conflict between tasks as the proportion of stimuli to be remapped. However, this definition does not take into account the nature of the relationship between competing outputs. Remapping a stimulus between similar responses (e.g., licking in the middle vs. to the left) is presumably easier than remapping between dissimilar responses with opposite behavioral meaning (e.g., licking vs. not licking). Thus, a more precise definition of task conflict should account for both input similarity and output dissimilarity. Future experimental designs that systematically vary output similarity while controlling input similarity—and vice versa—would provide suitable tools to disentangle their respective contributions to neural population modularity.

Beyond its role in handling conflicting sensorimotor mappings, modular population structure offers functional advantages during learning. Studies on ANNs have shown that adding a task-dependent gating signal, such that only sparse, mostly non-overlapping patterns of units are active for any one task, alleviates catastrophic forgetting [45, 46], especially when coupled with synaptic stabilization [47]. These authors further propose that the resulting modularity enables the separation into distinct neural populations implementing various cognitive processes, or building blocks, required to solve different tasks. Rather than learning an entirely new set of connection weights for each task, networks can leverage a task-dependent signal to recruit the necessary building blocks. This interpretation is supported by evidence that task representations in a recurrent network are compositional, emerging from the combination of representations of their building subtasks [3]. Therefore, in addition to improved robustness in the face of catastrophic forgetting, we propose that a modular structure, driven by task preference, provides a suitable substrate for transfer learning.

## 4 Methods

### 4.1 Dimension reduction

We chose to use dimension reduction techniques for two principal reasons: First, since the principal interest of this study was the representations of task variables, we targeted our analysis to the neural modes capturing most of inter-condition variance and get rid of the dimensions dominated by inter-trial variations. Furthermore, studying the geometry of these representations can prove challenging, as the probability of obtaining orthogonal axes by chance approaches one in high-dimensional spaces. To address this, we sought to conduct our tests in a low-dimensional space to ensure sufficient statistical power.

Unless otherwise mentioned, analyses focused on the average activity of neurons over a period of 1.5 seconds after stimulus presentation. We computed the average activity per condition for each animal and combined these average responses across animals to obtain a *c × n* matrix. *c* refers to the number of conditions (i.e. number of categories X number of choices) which was 10 across both tasks in the Response-switch paradigm (4 in GoNoGo and 6 in Left/Right). *n* corresponds to the total number of recorded neurons across animals (*n* = 2389). Next, the resulting time-averaged data matrix *M* was Z-scored for each column (i.e. for each neuron) so that 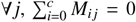 and 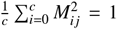. Principal Component Analysis (PCA) was then performed on *M* to identify which neural modes were most involved in inter-condition variations: *M*^*PCA*^ = *MW*, where *M*^*PCA*^ (*c × N*_*p*_) is the condition-averaged population activity projected on the top *N*_*p*_ principal components of *M*, and *W* is the PCA weight matrix (*n* × *N*_*p*_). To determine the number of component *N*_*p*_ to keep we compared the amount of variance explained by each component *k* to a distribution of variance explained by axes 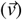 randomly drawn from Gaussian multivariate distribution and rescaled to unit norm.

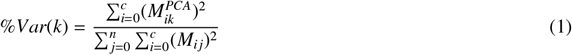

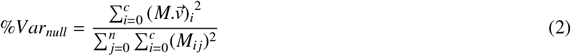

We considered that a component *k* captured significantly more variance than expected by chance if %*Var*(*k*) was above the 99.99 percentile of the random distribution 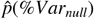, *p* < 10^−4^. We found that 9 out of 10 components explained significantly more inter-condition variance than expected by chance (Supplementary Figure 1a). Since the dimension of the inter-condition data was already small, we liberally kept these 9 components and only removed the last redundant dimension. This entire procedure was carried out before pseudo-population construction in order to obtain the same neural modes independently of the draws involved (see Pseudo-population) and improve the reproducibility of our results. Pseudo-population data was then projected onto these 9 components.

For a second part of the analyses we were interested in the temporal evolution of the activity within trials. Therefore, the space described above was inappropriate and another low-dimensional space was used. We computed mean neural time trajectories for each condition and a PCA model was trained on a (*c × t*) × *n* time data matrix. Where *t* stands for the number of time points in a trajectory (*t* = 18). As the number of sample (i.e. rows) in the condition-averaged trajectory matrix *T* was considerably larger this time, we favored a method inspired by cross-validated PCA (cvPCA[48, 49]) to establish the number of components to keep. The train-test covariance used in references ([48, 49]) did not decrease monotonically with component index in our dataset, making it difficult to define an appropriate cutoff. We therefore chose to determine the component cutoff based on the test-set explained variance, which decreased almost monotonically with component index (see Supplementary Fig. 1b). *T* was split 20 times into a train and test set using a two-fold procedure. The resulting matrices *T*_*train*_ and *T*_*test*_ (both 90 × 2389) were separately Z-scored for each column (i.e. neurons). PCA was fitted to the train set, 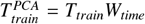, and test set was used to compute the mean explained variance by each of the components across splits 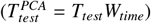.

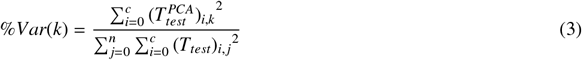

We then compared the amount of variance explained by each of the components to a null distribution computed from random axes as above.

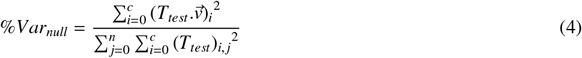

From this procedure, we determined that only the first 5 principal components met this criterion (see Supplementary Fig. 1b). As a consequence, neural trajectories from pseudo-population data were projected into the first 5 components of this second PCA model.

### 4.2 Geometry of population representations across individuals

In our preliminary analyses, we assessed angles between regression axes across individuals (Fig.2). The PCA weight matrices corresponding to each individual was obtained by decomposing *W* (see Dimension reduction) per block.

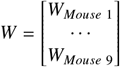

For each individual, the trial-wise data matrix *Y* was projected into its principal component space: 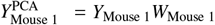. Following model fitting via ridge regression (see Ridge regression), we computed the angle between neural encoding axes corresponding to different pairs of regressors A and B. These axes were defined by their respective weight vectors in PC space, 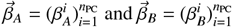. Angle between axes A and B was then computed as

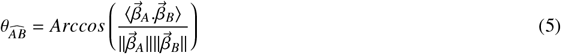

To assess the orthogonality of representation axes, we compared the observed angle between axes to a null distribution generated from random geometry. Specifically, we trained 1,000 models on regressor matrices with permuted rows, preserving the dimensionality and statistical properties of the original predictor-target pairs (X, Y) while disrupting their row-wise correspondence to obtain a reference distribution of inter-axis angles *p*(θ_null_). A one-sided Monte Carlo p-value was computed as *p* = 1 – percentile 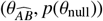, reflecting the likelihood of observing the measured angle under the null hypothesis of random orientation.

### 4.3 Pseudo-population

All neurons recorded from the nine mice were pooled together to construct a pseudo-population. We combined data from trials which shared the same experimental conditions across mice (i.e. same task, stimulus category, animal choice and licking response). Because of differences in performance the number of trials per condition was not the same for each animal. We therefore determined a number of trials that we aimed to incorporate for the pseudo-population that was neither too low to obtain a dataset capable of providing sufficient trials to fit the following models nor too high to avoid having to recycle trials from one animal. To do so, we used a conservative criterion: the number of trials per condition for the pseudo-population (referred as sampling level) was set to the third decile of the trial number distribution (see Supplementary Fig. 6). This way, only two animals were over-sampled, while seven were under-sampled. For each animal, we drew with replacement a number of trials per condition equal to the sampling level. Finally, within each condition, individual trial data randomly drawn were pooled across animals. At the end of the process, we obtained a *q × n* matrix, where *q* is the sum of sampling levels per condition (*q* = 381) and *n* is the sum of neurons for all nine animals (*n* = 2389). Using this approach, all the following analyses were found to be highly reproducible across different pseudo-population regardless of trial draws.

### 4.4 Ridge regression

#### 4.4.1 Motivation and framework

Due to the animals performing well in the task, stimulus category and animal choice were moderately correlated. Unregularized linear regression techniques are sensitive to these multi-collinearities by becoming highly variable from one iteration to the other, making it difficult to have a proper interpretation of the fitted estimator. As a consequence, we identified the neural modes encoding different task variables with Ridge regression. Ridge regression works with the advantage of not requiring unbiased estimators – rather, it adds bias to estimators to reduce the standard error. It adds bias enough to make the estimates a reliable representation of the population of data [50, 51].

The model we want to fit to the data is given by:

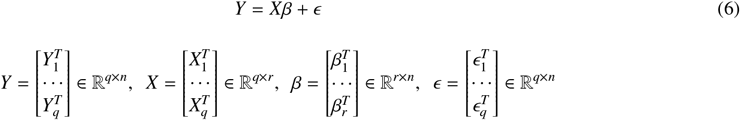

Where *q* corresponds to the number of pseudo trials, *n* the total number of neurons recorded across all animals and *r* the number of regressors considered.

The loss is thus defined as a L2-regularized residual sum of squares:

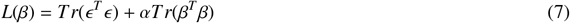

The scalar parameter *α* controls the level of regularisation over the regression weights *β*_*i,j*_ and *Tr* refers to matrix trace. The regression weights we ultimately used were those that minimised the loss function.

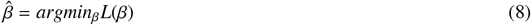

#### 4.4.2 Regressor matrix construction

To determine which regressors were relevant for our analyses, we first built a model that assumed task-independent representations, using single regressors for category and choice across the two tasks. Next, we compared its performance with a task-dependent model which assumed two distinct axes from one task to the other. We decomposed each regressor into a Go/NoGo part (forced to be null on L/R trials) and a L/R part (forced to be null on Go/NoGo trials). Using Akaike Information Criterion (AIC), we observed considerably better scores for the task-dependent model (Supplementary Fig. 5a,b), suggesting the need to split regressors from one task to the other to properly describe our data. These observations were in line with the very different category and choice axes finally obtained for one task versus the other (see Fig. 2j). In the end, we used a total of 7 different regressors to explain neural variability: Category Go/NoGo, Choice Go/NoGo, Category L/R, Choice L/R, Lick L/R, Task and Reward. Category refers to the category of the stimuli displayed whereas Choice refers to the animal choice, or reported category, in response to the stimuli. In the Go/NoGo task since the involvement of a motor response (i.e. licking the spout) is totally correlated with the animal choice we were not able to disambiguate between the two. However, it was not the case for the L/R task where the animal licking response was uncorrelated with animal reported category (Lick Left or Lick Right). As a consequence, Lick L/R refers to the regressor accounting for the motor response of the animal whereas Choice L/R the reported category (i.e the direction of licking). Finally, Task contrasts Go/NoGo trials versus L/R trials and Reward contrasts trials where the animal received water versus ones where it did not. A summary of the values of these regressors for each condition is given in a table (Supplementary Figure 5a).

We used an encoding scheme by trying to predict neural data merged across tasks and animals (*Y*) with the regressor matrix (*X*). First, neural data was denoised using PCA through transformation followed by back transformation to get rid of dimensions with inconsistent variations (see Dimension reduction). Afterwards, neural data was Z-scored. We next performed Grid Search to find the best regularization parameter value (*α*) over a 10 repeated 5-fold cross-validation procedure for 100 different *α* ranging exponentially from 10^−5^ to 10^4^. For each fold, to Z-score the regressor matrix, a standard scaler model was fitted to the train set and was run on the full dataset (train + test) to prevent information about the test set distribution to leak in the model during training. We kept the *α* value that gave the best mean test *R*^2^ scores for next steps (Supplementary Fig. 6c).

#### 4.4.3 cv*R*^2^ scores and statistics

To measure the amount of variance explained by each of the regressors, we computed cv*R*^2^ scores. To do so, we performed 10,000 permutations of regressor matrix (*X*) rows while keeping the order of data matrix (*Y*) rows. This broke the trial-to-trial correspondence between data and regressors, while maintaining the same statistics for both matrices. A first set of “fully permuted models” was obtained by permuting rows of *X* across all columns. In addition, for each column (i.e. regressor), we generated a set of “partially permuted models”. For each set, the rows of all columns were permuted in the same way as “fully permuted models”, except for the one column of the regressor of interest, which retained its structure. All permuted models were then fitted with the same pipeline as above, using a 5-fold cross-validation splitting strategy. On the one hand, *R*^2^ scores for partially permuted models gave us the amount of variance explained by the structure of a single regressor. On the other hand, *R*^2^ scores for fully permuted models gave us a null distribution that we would expect by chance, thanks to the statistics of the regressors. Finally, to correct for chance level, we calculated a cv*R*^2^ statistic for each permutation by subtracting the score of a fully permuted model from the score of its paired partially permuted model. For each regressor, we reported a cv*R*^2^ score that was taken as the average of this statistic across the 10,000 permutations. This procedure was repeated for each time point to obtain the temporal dynamics of the score for each regressor (Fig. 2i).

### 4.5 Analyses of neural selectivity

We interpreted regression weights as levels of selectivity for each regressor. Since we considered a total of seven regressors (see Ridge regression), each neuron was associated with a 7-dimensional selectivity vector. To quantify the extent to which a neuron was involved in the representations of variables in a given task, we computed the mean absolute selectivity across the task-specific regressors within that task (called “participation scores”). Since we could not disambiguate the animal’s choice with motor response in the Go/NoGo task (as they were completely correlated), some motor response contribution is dissimulated in the Choice GNG regressor. To avoid unfair comparison, we therefore also included the lick related activity in the LR task by incorporating the Lick LR regressor into the score. In this way we obtained participation scores with the same range of values across neurons in both tasks. Thus, participation scores in Go/NoGo (*P*_*GNG*_) and in Left/Right (*P*_*LR*_) were given by:

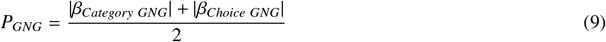

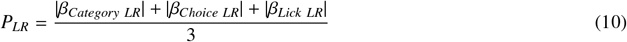

In addition, we compared how neurons mixed category and choice in each task. To quantify mixed selectivity, we computed the cosine similarity between 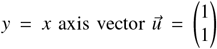 and selectivity vectors for variables in Go/NoG 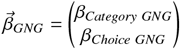 and in Left/Right 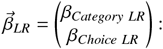

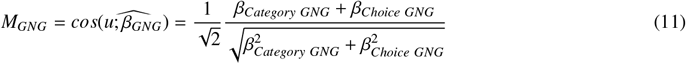

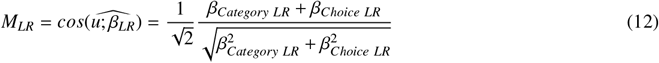

These two quantities were then plotted as a function of selectivity for the task (i.e *β*_*Task*_) in Figure 3. We observed a strong interaction between task selectivity, task participation and category-choice mixing. To ensure that this result is not due to some form of unanticipated circularity in our analyses we reproduced these results with two models trained over two different time periods. The selectivity for the task was obtained by performing a regression on the mean activity over a period of 1.5s pre-stimulus. In contrast, selectivities to Category Choice and Lick response were obtained from a model trained on mean activity over a period of 1.5s post-stimulus.

### 4.6 Subpopulations and Gaussian Mixture

#### 4.6.1 Selectivity space

After linear regression, each neuron was associated with a selectivity vector *s*_*i*_ composed of the weights for each of the seven regressors. The information about the selectivity of all the neurons is summarized in a matrix

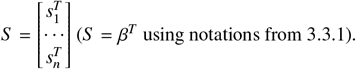

#### 4.6.2 ePAIRS

We first sought to quantify the amount of local structure in the selectivity space using the ePAIRS test. ePAIRS is a non-parametric test that aims to quantify neural selectivities clustering across the population. To do so, we examine the distribution of selectivity vector directions and compare it to a null distribution generated by a multivariate Gaussian with a covariance matrix identical to the covariance of our data. A significant outcome of the ePAIRS test indicates that the empirical distribution presents multiple “preferred” directions which are incompatible with a random Gaussian structure. In detail, we followed these steps:

1. *S* was centered around 0 by removing the mean for each column, so that for each 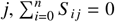.
2. For each selectivity vector *s*_*i*_, we determine its *k* nearest neighbors using the cosine metric. That is to say, we consider as neighbors the *k* vectors for which the quantity 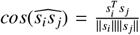 is the highest. With *k* an hyperparameter set to 2 in our analysis.
3. For each selectivity vector, we compute the mean angle *θ*_*i*_ with its *k* nearest neighbors, defining an empirical distribution 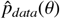.
4. Next, a multivariate Gaussian distribution *N* (0, Σ) is fit to the set of selectivity vectors *S*, with Σ the empirical covariance of *S*, computed as 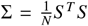. Then, we applied steps 2 and 3 on 200 samples of the multivariate Gaussian with the same number *n* = 2389 of vectors. Angles from these 200 samples are then concatenated to obtain a null distribution 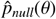.
5. To assess the statistical significance of the difference between the data and the null distribution we computed the PAIRS statistic

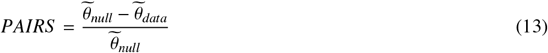

Where 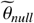 and 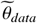 represents the median of 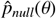 and 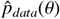 respectively. This statistic is 1 if all neurons have at least *k* identical partners, and 0 if clustering is only as strong as expected by chance. Since we could compute 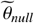 for each of the 200 samples separately, we used these values to compute PAIRS statistic for every pair of random samples and find a null distribution of the PAIRS statistic expected by chance. A one-sided non-parametric p-value was then computed by taking 1 minus the percentile of the empirical PAIRS statistic in this null distribution.
6. Finally, the magnitude of the effect is computed as a Cohen’s effect size *d*

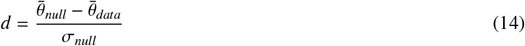

Where 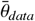 represents the mean of 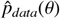while 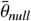 and 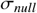 represent respectively the mean and the standard deviation of 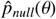. A positive effect size *d* > 0 indicates that angles between neighbors are smaller in the data compared to the random Gaussian distribution, meaning that selectivity vectors are more highly clustered than expected. On the contrary a negative effect size *d* < 0 indicates that vectors are more regularly dispersed than expected from random.

#### 4.6.3 Gaussian mixture

Since we found that the distribution of selectivities was not random, we wanted to know if we could identify subpopulations with similar selectivity properties. This was done using a Gaussian Mixture (GM) algorithm that was fit to describe all 7-dimensional selectivity vectors by a combination of 7-dimensional Gaussian distributions. We used the “Bayesian-GaussianMixture” model from scikit-learn package. The model was trained on the selectivity matrix *β* with 20 unique initializations (*n*_*init*_ = 20). Just like for our PCA model (see Dimension reduction), we used a cross-validated approach for GM models (cvGM) to determine the optimal number of components to describe selectivities. GM models with a number of components varying from 1 to 10 were trained on half of selectivity vectors and tested on the other half 100 times using a two-fold procedure on selectivity matrix. For each split and each model, test scores were computed as the per-sample average log-likelihood of the test set. The score increase associated with the addition of the *n*th component was calculated as the mean score difference between the *n*-components model and the (*n ™* 1)-components model across splits. This way, we reliably obtained a positive and monotonically decreasing score increase as more components were added. The number of components retained was the last for which the score increase was still statistically significantly greater than 0. This criterion was met if the score increase was positive for more than 95% of the splits (*p* < 0.05). Finally, we used the density of mixture components for each neuron to determine which subpopulation it belonged to. We considered a neuron to belong to population X if the density of component X was greater than 0.9. Thus, about 80% of the neurons belonged to a single subpopulation, while the remaining part could not be assigned to any subpopulation and was excluded from the analyses.

### 4.7 Recurrent neural network modelling

#### 4.7.1 Description

To investigate the mechanisms involved in the emergence of task-specialized subpopulations in our dataset, we developed an artificial recurrent neural network. The network was composed of the same number of units *N* as the dataset (*N* = 2389). Connectivity between units was taken as random by obtaining a weight matrix *W* with independent draws from Gaussian distribution 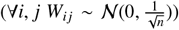. Each unit *i* received a total input *x*_*i*_ composed of inputs from other units and external inputs. The total input was described for all neurons by the vector 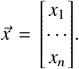 The network receives *x*_1_ three kind of external inputs :

1. An external input reflecting stimulus category, *s*(*t*) = ±1 + η_*s*_(*t*) with ∀*t*, η_*s*_(*t*) ∼ *N* (0, 1). Stimulus category input was fed differently to each unit based on a set of weight 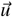 with ∀*i, u*_*i*_ ∼ *N* (0, *σ*_*stimulus*_).
2. Similarly, a task input *c*(*t*) = ±1 +η_*c*_(*t*) was fed to the network units via 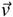 with ∀*i, v*_*i*_ ∼ *N* (0, *σ*_*task*_). ∀*t*, η_*c*_(*t*) ∼ *N* (0, 1).
3. A constant modulation, independent of task conditions, which could be either globally excitatory or inhibitory across the population. This global modulation input was fed to the network via 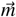 with ∀*i, m*_*i*_ ∼ *N* (*µ*_*modulation*_, 1).
4. An external noise 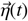 independently drawn for each unit and at each time point from gaussian distribution with ∀*i*, η_*i*_ ∼ *N* (0, 5).

The three parameters *σ*_*stimulus*_,*σ*_*task*_ and *µ*_*modulation*_ were the ones used to fit the model to the data.

Total input was converted to an activity level 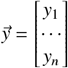. by applying a sygmoidal non-linearity

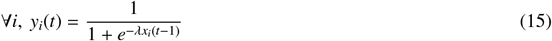

Where *λ* = 3 is a scaling factor that represents the steepness of the sygmoid. Additionally, the network dynamic followed a first order differential equation.

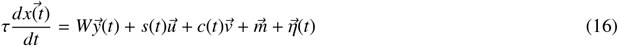

This equation was numerically approximated using Euler’s method over 30 iterations for each trial.

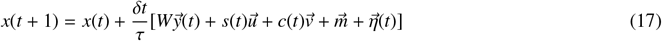

Where *τ* is the characteristic time of units activities and *δt* is the simulation time step taken as 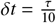.

#### 4.7.2 Model fitting

To fit the model we performed a grid search over the set of parameters {*σ*_*stimulus*_, *σ*_*task*_, *µ*_*modulation*_}. In detail, we followed these steps:

1. We built models for parameter values linearly ranging from 0.5 to 1.5 for *σ*_*stimulus*_ from 1 to 5 for *σ*_*task*_ and from −3 to 3 for *µ*_*modulation*_ (8 points for each, i.e. a total of 8^3^ = 512 models).
2. Each model was run on *p* = 100 trials with an homogeneous distribution of conditions (25 trials for two stimulus categories in two tasks).
3. In the same way as in the dataset, units activity were averaged over time within each trial resulting for each model in a *p* × *N* simulated data matrix *Y*.
4. We then performed Ridge regression on *Y* using two task-specific category regressors: (‘Category GNG’,’Category LR’) and a ‘Task’ regressor.
5. We computed simplified participation scores (see Analyses of neural selectivity) for each task *P*_*Task*_ = |*β*_*Category Task*_| and averaged these scores across units according to their task selectivity to obtain two simulated distributions 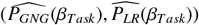 as in the data.
6. Finally, models performance in reproducing data were evaluated with the *R*^2^ score:

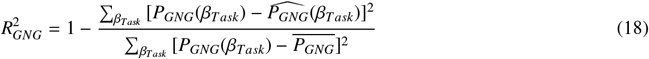

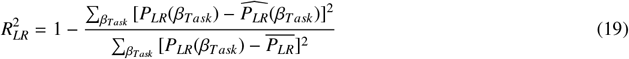

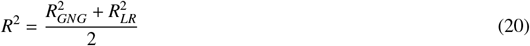

Where 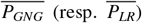 corresponds to the mean participation scores in Go/NoGo (resp. Left/Right) across task selectivities for the distribution observed in the data. The model maximizing this score was kept for further analyses.

#### 4.7.3 ePAIRS and Gaussian Mixture on simulated units

Model units were associated with a selectivity vector based on the regression weights calculated during model fitting. To better understand how contextual input and global modulation interact to give rise to functional subpopulations we performed ePAIRS tests on several models with the same procedure as for the data (see ePAIRS). To obtain better resolution in the plane of the parameter of interest {*σ*_*task*_, *µ*_*modulation*_}we set the stimulus signal to the value giving the best fit to the data (*σ*_*stimulus*_ = 1). We then generated models for values linearly ranging from 1 to 5 for *σ*_*context*_ and from −3 to 3 for *µ*_*modulation*_ (20 points for each, i.e. a total of 20^2^ = 400 models). We reported effect sizes obtained for each model with a surface plot smoothed by a Gaussian kernel with a standard deviation of *σ*_*Kernel*_ = 2.

To ensure that the clustering of selectivities captured by the ePAIRS tests was indeed of the same nature as in the data, we further characterized it using a Gaussian mixture algorithm. We determined the number of components needed to describe selectivity distribution and ran the GM model using the same pipeline as for the data (see Gaussian mixture).

#### 4.7.4 Model performance on the two tasks

We assessed the performance of the best-fit model on both Go/NoGo and Left/Right tasks. We did so by training two single layer perceptron readouts (from scikit-learn package). Each perceptron was fed with post-stimulus units’ mean activity. A first ‘Lick’ readout was trained to output 1 if the animal had to lick in the given trial and −1 otherwise. A second ‘Tongue direction’ readout was trained to output 0 if the animal had to lick a center spout (for Go/NoGo task) −1 if it was a left spout and +1 if it was a right spout (Left/Right task). The correct ‘Lick’ and ‘Tongue direction’ responses were determined based on the true stimulus category sent to the network, using the same task design as for the data. Readouts were trained with stochastic gradient descent on half of the simulated trials and tested on the other half, using a 10 repeated 2-fold cross-validation procedure. Accuracy was computed across folds and for each task as the percentage of trials from test set in which both readouts answered correctly.

## 5 Supplementary information

**Supplementary Figure 1.**
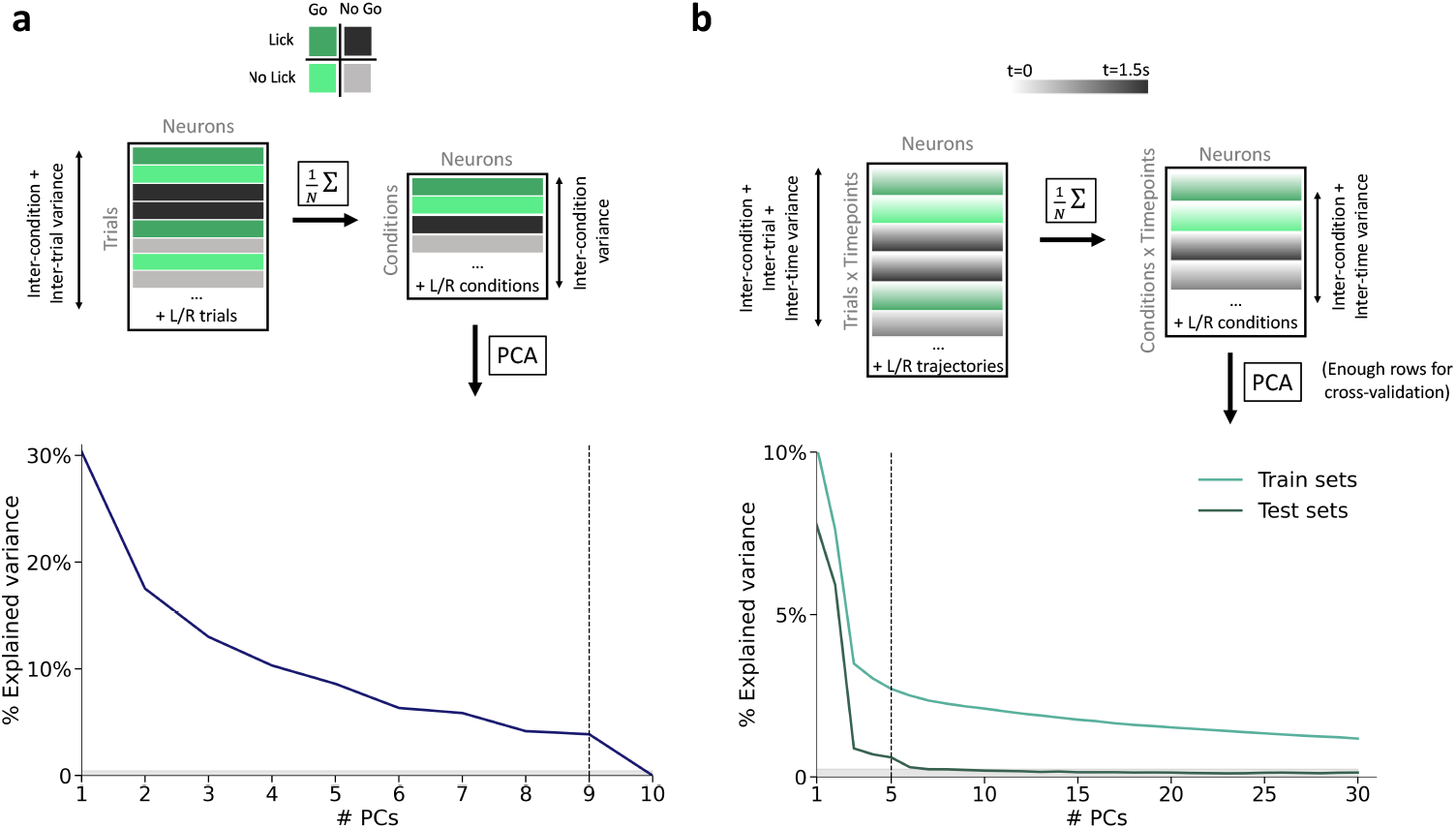
**a**, Training procedure and explained variance for PCA model trained on time-averaged data. Top, initial data-matrix was reduced to a condition-averaged data matrix that highlights inter-condition variations. Bottom, inter-condition variance explained by each PCs. Grey shaded area corresponds to explained variance distribution expected by chance (99.99 percentile). Vertical dotted line corresponds to the PC cuttoff (line included) used in the analysis. **b**, Same as **a** for time trajectories. Top, trajectory data-matrix was averaged for each condition to highlight inter-time and inter-condition variations. Because of the number of time points (*t* > 18) the resulting matrix had considerably more rows enabling us to carry out proper cross-validation. Bottom, train and test sets variance explained by each PCs. Grey shaded area corresponds to the test set explained variance distribution expected by chance (99.99 percentile). Vertical dotted line corresponds to the PC cuttoff (line included) used in the analysis.

**Supplementary Figure 2.**
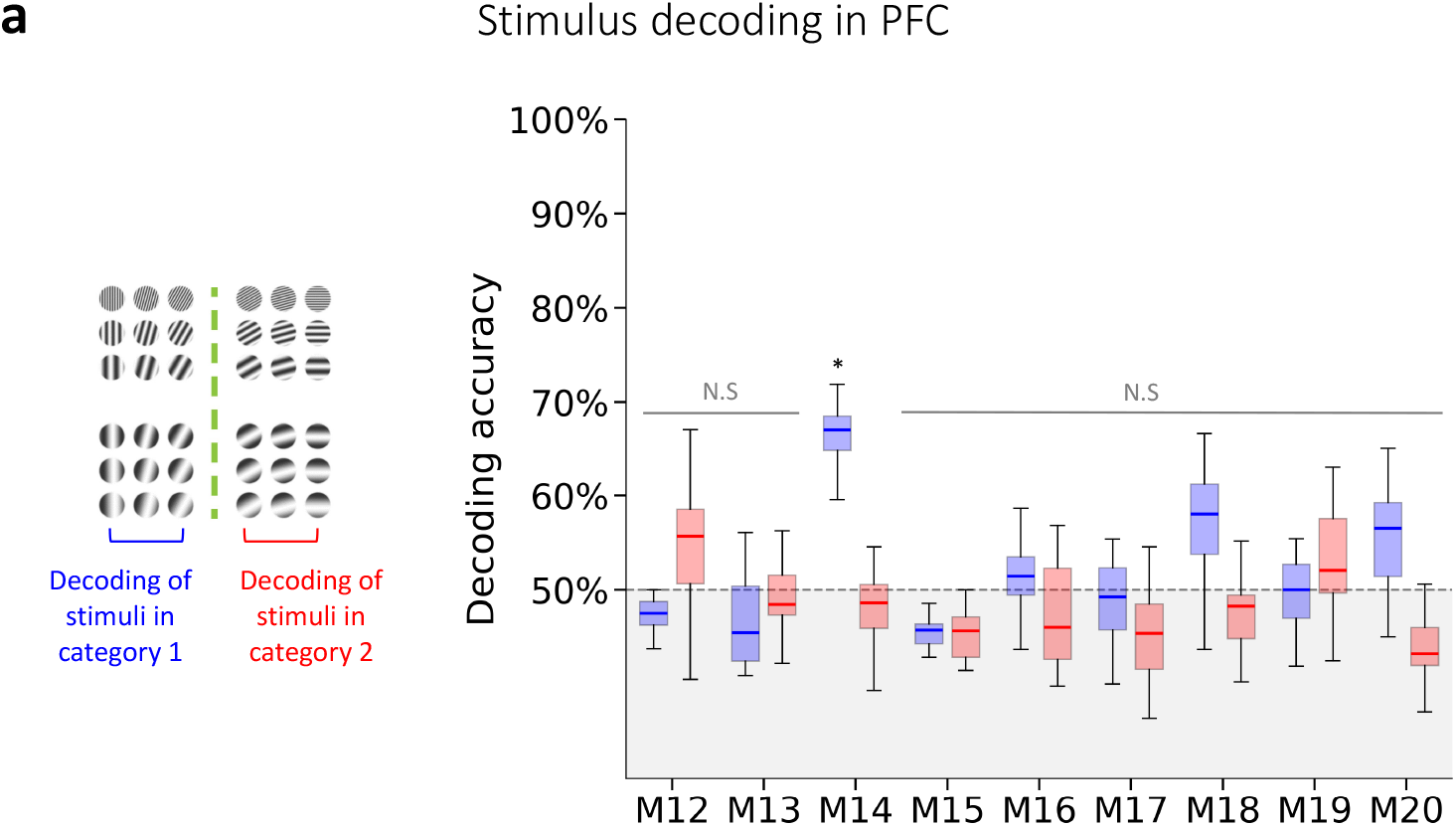
**a**, Linear decoding of stimulus identity within a category across animals from mPFC population activity. Blue, SVC decoder performance for decoding stimulus from category 1. Red, decoding performance for stimulus from category 2. Decoders were trained using a 5-fold cross-validation procedure. Significant decoding performances were assessed with permutation tests corrected for multiple comparisons, Bonferroni-corrected (N.S, not significant, * *p* < 0.05).

**Supplementary Figure 3.**
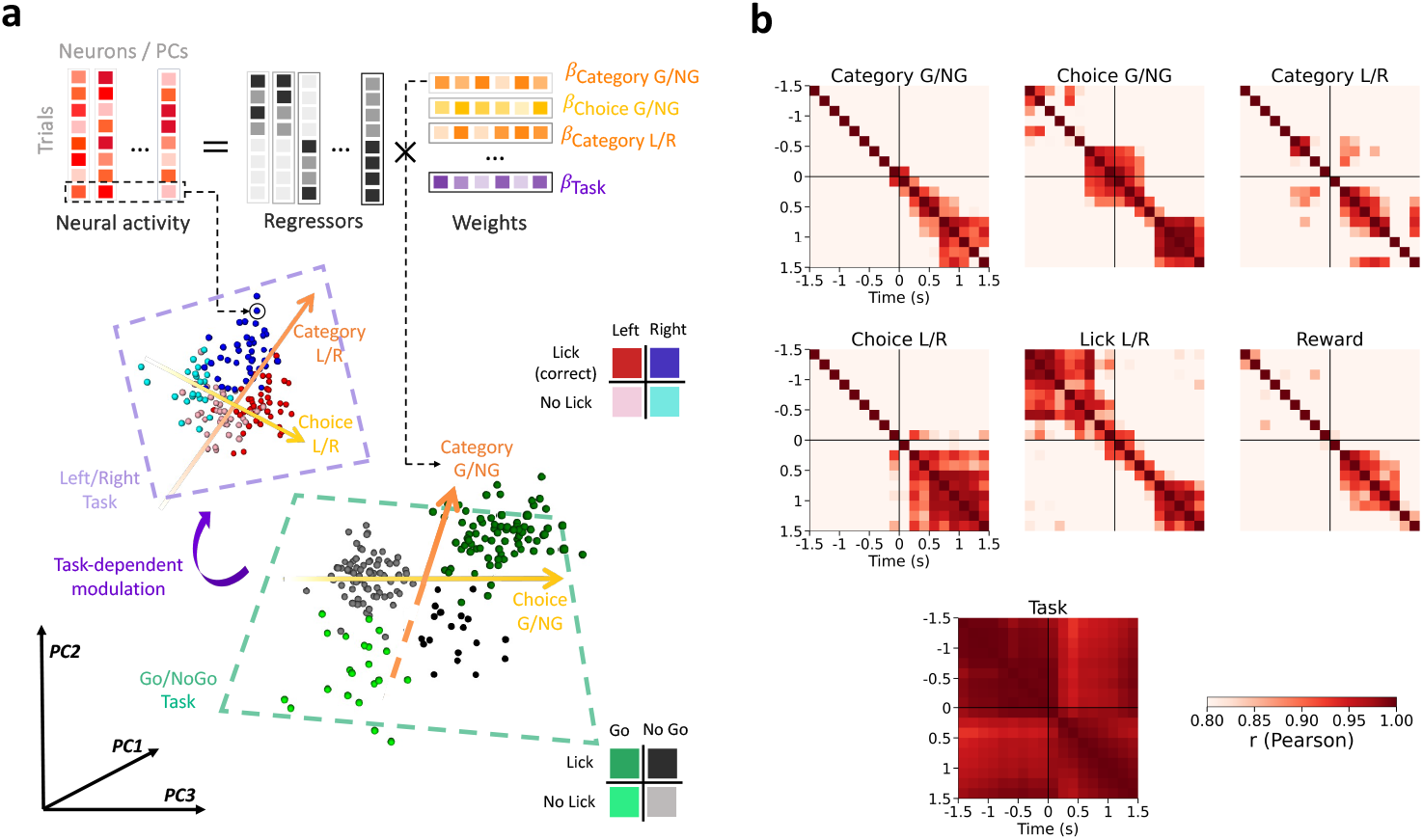
**a**, Top, linear regression of neural activity across the two tasks using multiple behaviorally-relevant regressors (G/NG:Go/NoGo, L/R:Left/Right). Bottom, pseudo-population vectors for every pseudo-trials projected on a subset of first principal components. Pseudo-trial labels are indicated by top-right color code (for Left/Right task) and bottom-right color code (for Go/NoGo task). Dotted 2D plans correspond to subspaces that explain most of the variance in a given task. Orange and yellow arrows correspond respectively to encoding axes for the category of the stimuli and the choice of the animal. **b**, Cross-correlation through time of neural axes encoding for each regressor (*t* = 0 corresponds to stimulus onset). For each regressor, only models with a significant explained variance (*cvR*^2^) were considered to compute cross-correlation.

**Supplementary Figure 4.**
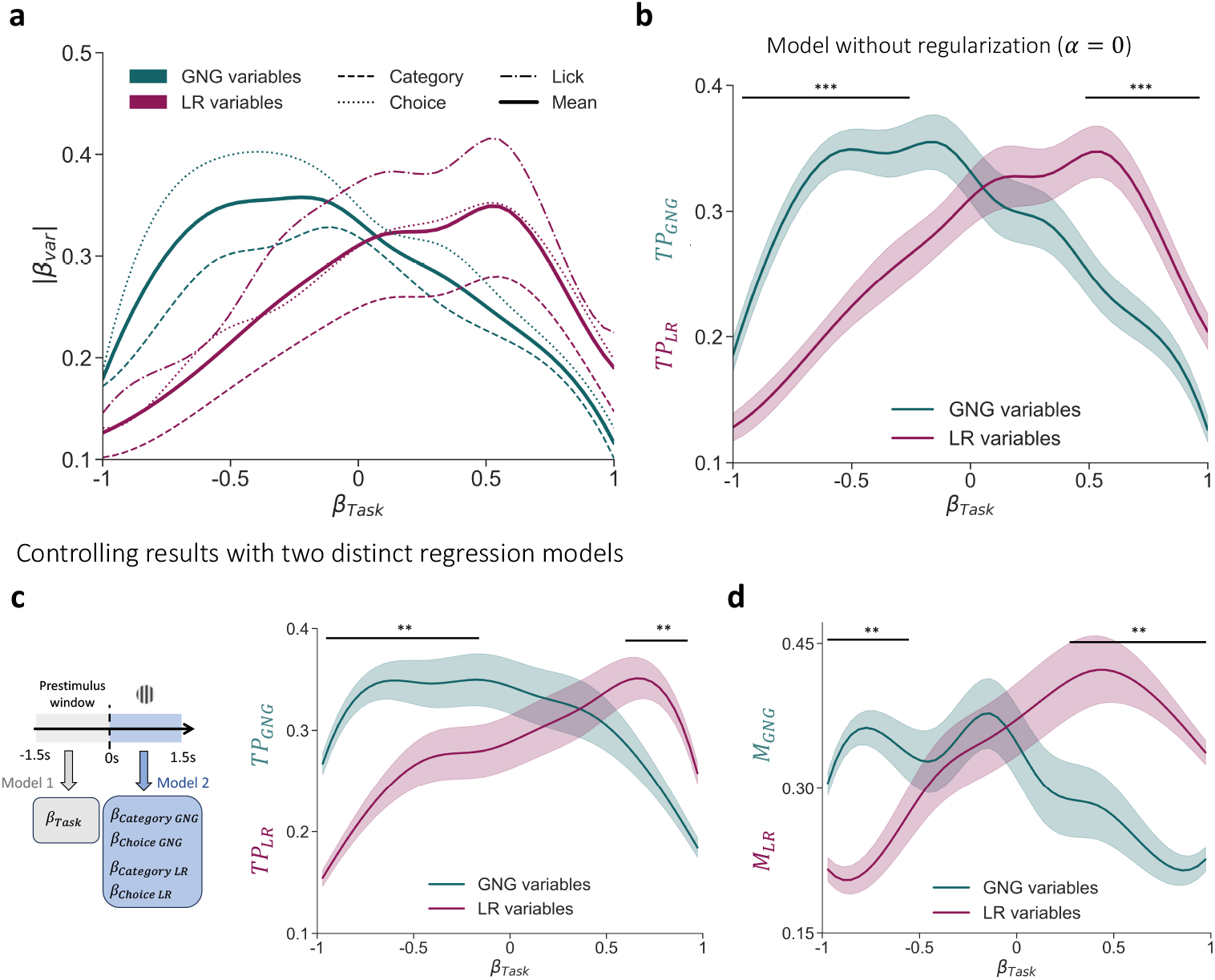
**a**, Decomposition of each selectivity in task participation score. Absolute selectivity to the variables from Go/NoGo (Choice and Category) are represented in dark green. For Left/Right task, the animal choice (i.e. reported category) and its motor responses (i.e. licks) were decorrelated and we were able to further disambiguate their contribution in the regression. **b**, Task participation scores reproduced with a linear model trained without regularization. Regularization had little effect on this result.**c**, Reproduction of task participation scores from Fig 3b using two distinct linear regression models. Selectivity for the task was obtained from averaged activity on pre-stimulus window while selectivity for task variables (Category, Choice) were obtained from post-stimulus activity. **d**, Same as in **c** to reproduce mixing scores from Fig. 3c. Despite quantitative disparities, the main effects reported in Fig. 3 remained present with this control.

**Supplementary Figure 5.**
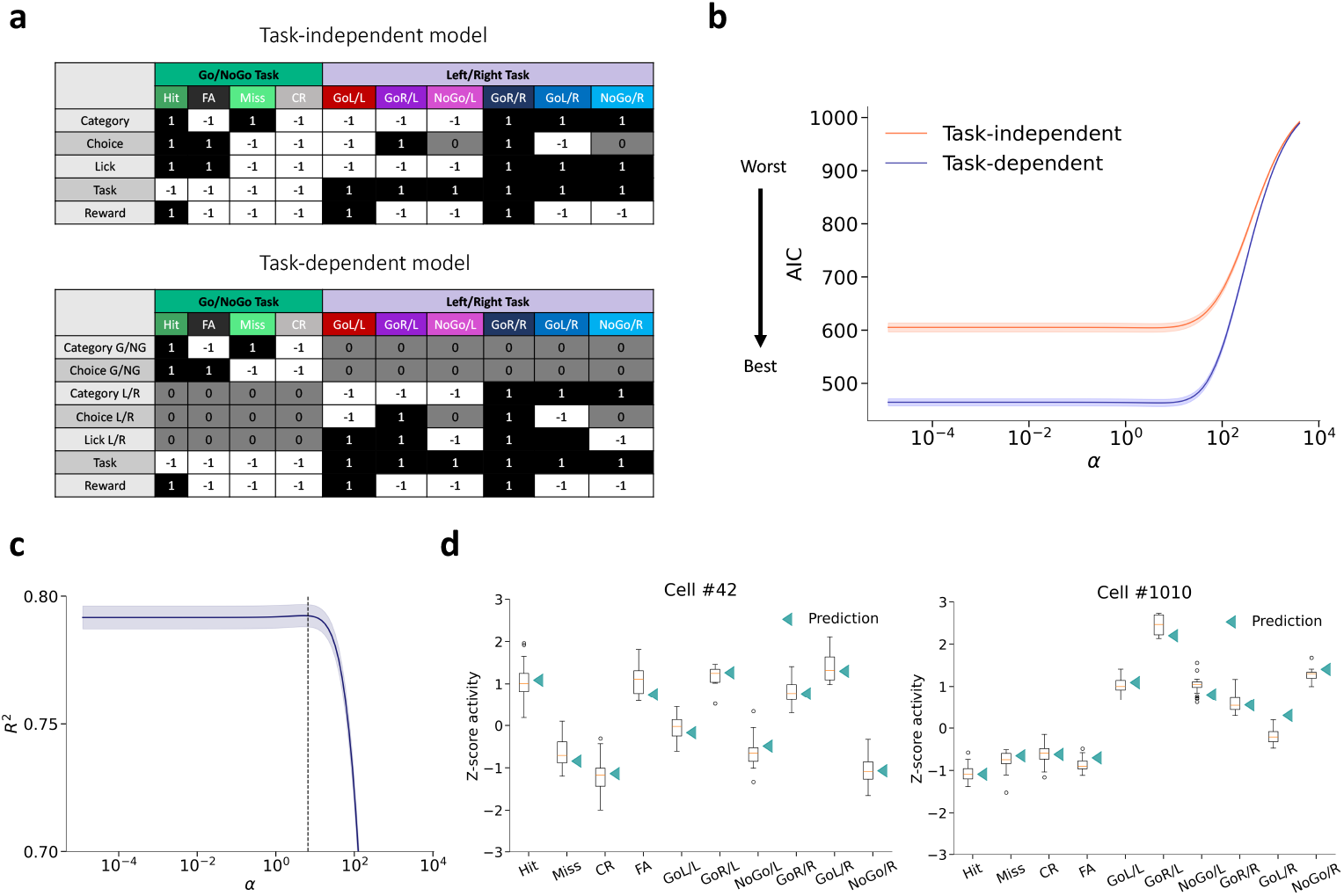
**a**, Design matrices for task-independent (top) or task-dependent (bottom) representations of category and choice in population activity. GoR (resp GoL) : the animal licked to the right (resp left), /R (resp /L) : stimulus category implied to lick to the right (resp left) to be rewarded. **b**, Akaiki information criterion (AIC) for the two models across different regularization coefficients (*α*). Note that the blue line is below the orange one for the range of *α* retained for the analysis (*α* < 10, see **c**) meaning that task-dependent decomposition of regressors is justified, following Akaike criterion, as it yields significant improvement in model performance. **c**, Mean *R*^2^ score across folds in predicting data matrix from test set trials. *R*^2^ is plotted for a range of regularization coefficients. Vertical dotted line corresponds to *α* that provided best cross-validation scores and was retained to fit the model to the data. **d**, Prediction of the activity of 2 example cells across trials from different conditions. White box-plot corresponds to Z-scored activity distribution from test set trials of a given condition.

**Supplementary Figure 6.**
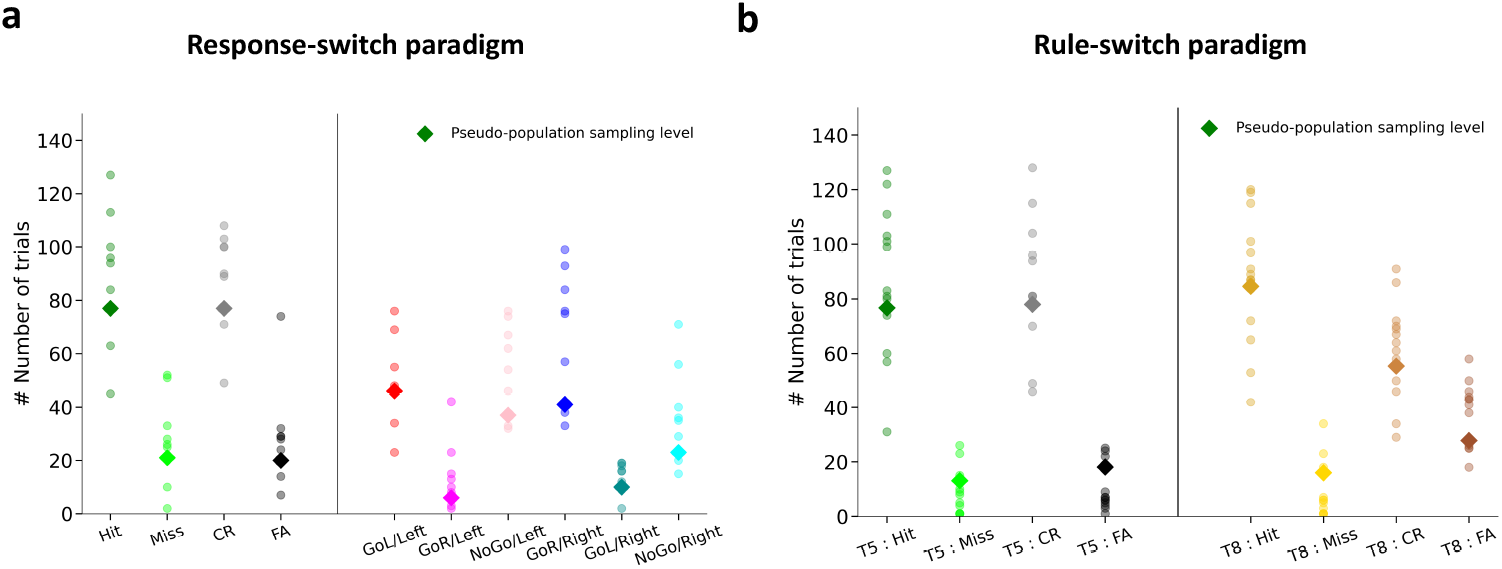
**a**, Number of trials of each condition from Response-switch across all individuals (circles). Diamonds corresponds to sampling level used to construct pseudo-population (third decile of number of trials distribution). For each mouse and each condition a number of trials equal to the sampling level was pooled with replacement. Neurons recorded from all mice were pooled together to obtain a number of pseudo-trials equal to the sampling level for each condition. **b**, Same as **a** for Rule-switch paradigm. In this dataset, Misses and FAs were considerably under-represented and sampling level was corrected (fifth decile) to obtain enough pseudo-trials.

## Acknowledgements

We thank Joao Barbosa and Célian Bimbard for careful reading of the manuscript and precious comments.

## Data availability

Data are available in the initial study by Reinert et al 2021 [11].

## Code availability

The analysis scripts are freely available at https://github.com/HTissot42/Sensorimotor-remapping-drives-task-specialization

## Author contribution

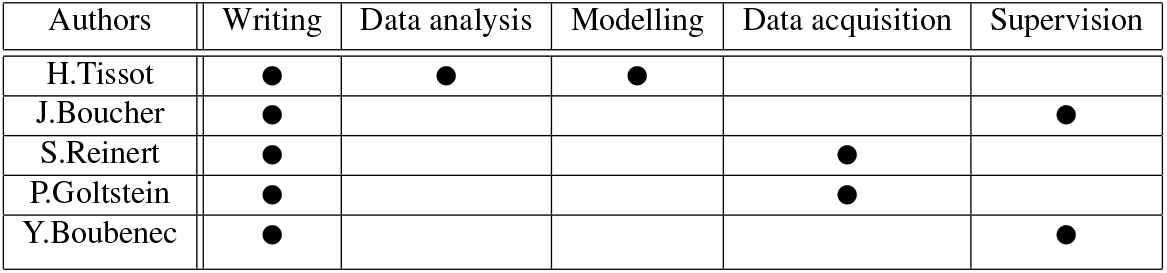

